# hENT Inhibition Prevents Pyrimidine-Driven Resistance to DHODH Inhibition in Malignant Rhabdoid Tumors

**DOI:** 10.64898/2026.01.25.701565

**Authors:** Marjolein M.G. Kes, Giulia Perticari, Jolanda Kooiman, Nadia Anderson, Jeroen W.A. Jansen, Esther A. Zaal, Celia R. Berkers, Jarno Drost

**Affiliations:** Princess Máxima Center for Pediatric Oncology, Utrecht, The Netherlands; Oncode Institute, Utrecht, The Netherlands; Division Cell Biology, Metabolism & Cancer, Department Biomolecular Health Sciences, Faculty of Veterinary Medicine, Utrecht University, Utrecht, The Netherlands

## Abstract

Extracranial malignant rhabdoid tumors (ecMRTs) are aggressive pediatric cancers characterized by mutations in genes encoding members of the SWItch/Sucrose Non-Fermentable (SWI/SNF) chromatin remodeling complex, with limited effective treatment options. Metabolic reprogramming, including nucleotide biosynthesis, is a hallmark of cancer that represents a potential therapeutic vulnerability. Previously, we demonstrated that inhibition of the *de novo* pyrimidine synthesis enzyme dihydroorotate dehydrogenase (DHODH) represents a promising treatment strategy for rhabdoid tumors, including ecMRTs and atypical teratoid/rhabdoid tumors (AT/RTs). Using patient-derived tumoroid models, we now extend these findings by showing that another SWI/SNF-deficient pediatric cancer, small cell carcinoma of the ovary, hypercalcemic type (SCCOHT), is also sensitive to DHODH inhibition. Gene expression analyses further confirmed that SCCOHT and AT/RT, like ecMRT, exhibit elevated expression of *de novo* nucleotide synthesis genes. To determine whether metabolic features of the tumor microenvironment (TME) influence DHODH inhibitor (DHODHi) response, we profiled plasma and tumor interstitial fluid from orthotopic ecMRT-bearing mice and found that the TME was markedly enriched in nucleosides and nucleobases. Supplementation of standard culture media with these nucleosides demonstrated that pyrimidines, but not purines, can rescue ecMRT cells from DHODHi-induced growth inhibition via activation of the pyrimidine salvage pathway. This resistance mechanism was effectively reversed by co-treatment with human equilibrative nucleoside transporter (hENT) inhibitors, enhancing DHODHi sensitivity in ecMRTs. Collectively, these findings reveal a conserved metabolic vulnerability across SWI/SNF-deficient pediatric cancers and emphasize the critical importance of modeling tumor metabolism under physiologically relevant conditions to accurately identify metabolism-targeted therapeutic strategies for these currently lethal pediatric cancer types.

## Introduction

Malignant rhabdoid tumors (MRTs) are rare and highly aggressive embryonal neoplasms that mainly affect children under three years of age^1^. These tumors can develop in multiple anatomical locations, with the central nervous system (CNS) being the most common site^2^. Within the CNS, MRTs are referred to as atypical teratoid/rhabdoid tumors (AT/RTs)^2^. Extracranial MRTs (ecMRTs), in contrast, can occur in various soft tissues and visceral organs, such as the liver, bladder, retroperitoneum, extremities, and pelvis, as well as the kidney^3^. MRT pathogenesis is driven by aberrant differentiation and lineage specification during fetal development, caused by the biallelic inactivation of either *SMARCB1* (∼95% of cases) or *SMARCA4* (∼5% of cases)^4–7^. These genes encode core subunits of the SWItch/Sucrose Non-Fermentable (SWI/SNF) chromatin-remodeling complex, which plays a critical role in transcriptional regulation by nucleosome repositioning^8–10^. Current MRT treatment strategies rely on multimodal therapy, including chemotherapy, radiotherapy, and surgical resection^11^. However, prognosis remains extremely poor, particularly in younger patients and those with metastatic disease^12^, underscoring the urgent need for novel therapeutic approaches.

Elevated nucleotide synthesis and utilization are recognized as fundamental metabolic dependencies in cancer cells, irrespective of tumor type and genetic background^13^. This dependency arises because nucleotides are rate-limiting for key processes that are hyperactive in cancer cells, including DNA replication and repair^14^, transcription^15^, ribosome biogenesis^16^, and differentiation blocking^17^. To exploit this vulnerability, nucleotide metabolism-targeting chemotherapies have been a cornerstone of cancer treatment for decades^18–21^. Classic antimetabolites such as methotrexate (MTX)^18,19^ and 6-mercaptopurine (6-MP)^20,21^ have not only shaped modern oncology but also provided foundational insights into the metabolic dependencies of cancer cells.

Nucleotides are classified into two main groups: purines and pyrimidines. Both are synthesized via two principal routes: the *de novo* synthesis and the salvage pathways. The *de novo* synthesis pathway is an energy-intensive process that assembles nucleotides from small precursors, while the salvage pathway efficiently recycles preformed nucleosides or nucleobases into nucleoside monophosphates (NMPs) through a single enzymatic step. Traditionally, highly proliferative cells were thought to rely primarily on *de novo* nucleotide synthesis, whereas differentiated tissues were thought to predominantly utilize the salvage pathway^13^. In line with this assumption, our previous work^22^ demonstrated MRTs exhibit pronounced activation of the *de novo* nucleotide synthesis pathway. Additionally, patient-derived ecMRT and AT/RT tumor organoids (tumoroids) showed an increased sensitivity to BAY-2402234 (BAY), an inhibitor of the *de novo* pyrimidine biosynthesis enzyme dihydroorotate dehydrogenase (DHODH), compared to normal kidney organoids^22^. However, it is unknown whether this metabolic vulnerability extends beyond ecMRTs and AT/RTs to additional pediatric malignancies driven by mutations in SWI/SNF complex components, such as small cell carcinomas of the ovary, hypercalcemic type (SCCOHTs), characterized by recurrent loss of *SMARCA4* in the vast majority of cases^23–25^. Moreover, conventional cell culture conditions lack the nucleosides and nucleobases typically present in the tumor microenvironment (TME), leading to an increased reliance on *de novo* nucleotide synthesis and possibly an artificially enhanced sensitivity to DHODHi *in vitro*. Consequently, the relative contributions of the *de novo* synthesis and salvage pathways to nucleotide metabolism in MRTs under physiologically relevant conditions remain poorly understood, representing a critical gap in accurately modeling and targeting this pediatric cancer type, and cancer in general.

Using patient-derived tumoroid models, we evaluated the sensitivity of ecMRTs to a panel of DHODHi and extended our investigation to SCCOHTs to assess whether this dependence on *de novo* nucleotide biosynthesis constitutes a shared metabolic vulnerability across different SWI/SNF-deficient pediatric cancers. To better capture the *in vivo* metabolic context, we profiled plasma and tumor interstitial fluid (TIF) from orthotopically implanted ecMRT-bearing mice. This approach revealed tumor microenvironment-derived nutrients that drive resistance to DHODHi and enabled the identification of rational combination strategies to prevent this resistance – insights not achievable using standard *in vitro* culture conditions.

## Results

### MRTs display selective sensitivity to a wide range of DHODH inhibitors

In our previous work (Kes *et al*., 2025^22^), we identified BAY-2402234 (BAY) as a promising treatment strategy for extracranial malignant rhabdoid tumor (ecMRT) and atypical teratoid/ rhabdoid tumors (AT/RT). BAY is a well-characterized inhibitor of dihydroorotate dehydrogenase (DHODH), a key enzyme in the *de novo* pyrimidine synthesis pathway (**Figure 1A**). Inhibition of DHODH with BAY indeed lowered nucleotide levels and significantly induced apoptosis in patient-derived ecMRT tumoroids^22^. While these findings highlight the therapeutic potential of DHODH blockade in MRT, BAY is not currently approved for clinical use. To identify additional clinically actionable candidates for further (pre-)clinical development, we sought to evaluate alternative DHODH inhibitors (DHODHi) with advanced development in oncology^37^, specifically AG-636, Farudodstat (ASLAN003), PTC299, and GTX-196.

**Figure 1.**
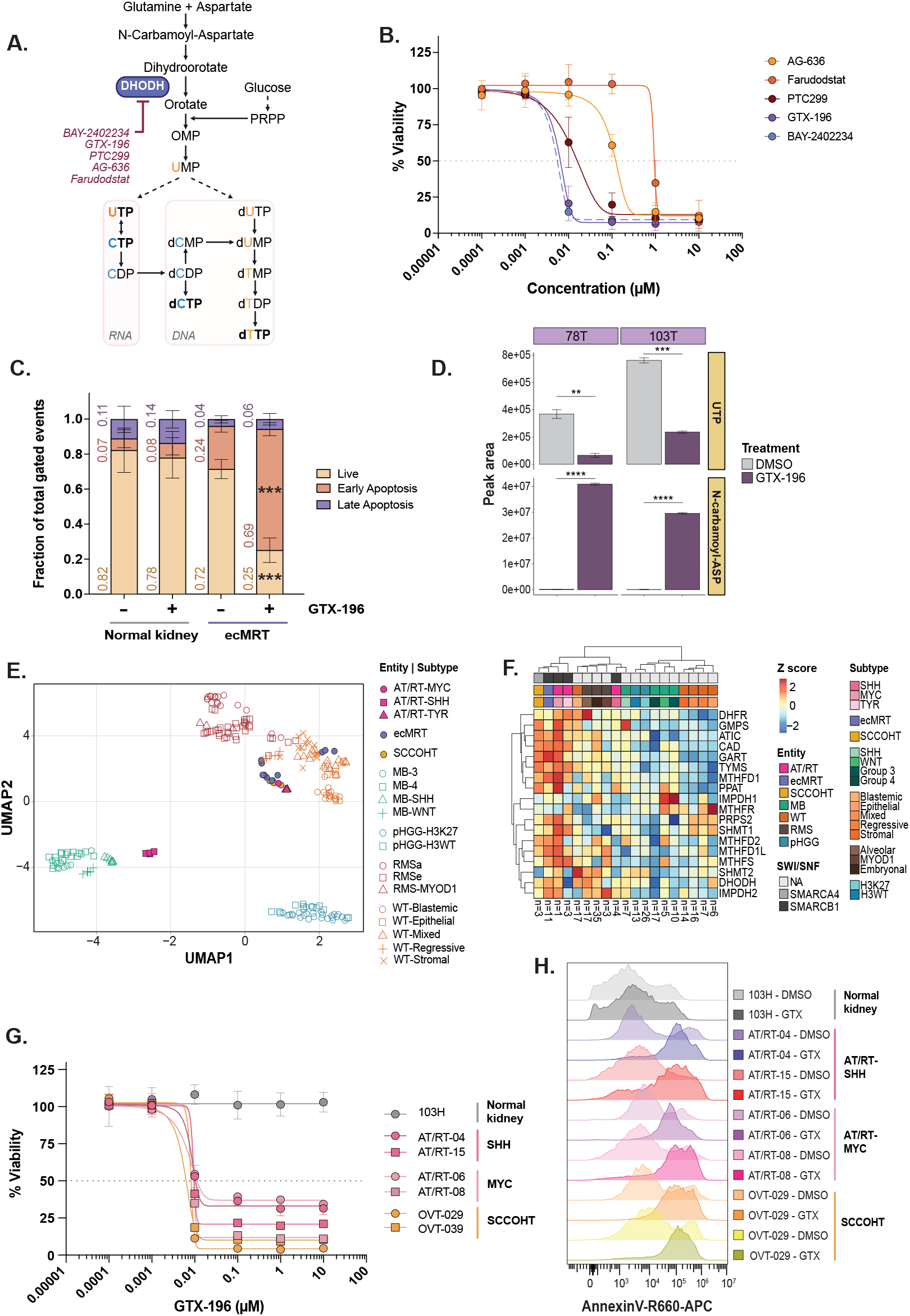
Inhibition of *de novo* pyrimidine synthesis enzyme DHODH marks a vulnerability of SWI/SNF-mutant tumors. **A)** Simplified schematic of the *de novo* pyrimidine biosynthesis pathway, depicting key enzymatic steps leading to the formation of nucleotide triphosphates. The final products – UTP, CTP, dCTP, and dTTP – used as direct substrates for RNA and DNA synthesis are indicated in bold. The key enzyme dihydroorotate dehydrogenase (DHODH) is highlighted in purple and shown alongside DHODH inhibitors (DHODHi) with relevance in oncology. **B)** Dose-response curves of DHODHi for patient-derived ecMRT tumoroid cultures. Data points are represented as the mean ± SD of three independent ecMRT models, with each model measured in quadruplicate. Data are normalized to DMSO vehicle (100%). The grey dotted horizontal line represents a viability of 50% (IC_50_). **C)** Bar graphs representing live (yellow), early apoptotic (orange), and late apoptotic (purple) cell fractions of ecMRT tumoroids and normal kidney organoids upon treatment with 50 nM GTX-196 or DMSO vehicle control for 120 h. Data are represented as the mean ± SD of *n = 3* independent models. **D)** Bar graphs depicting the total peak area, corresponding to metabolite levels of uridine diphosphate (UDP) and N-carbamoyl-aspartate in ecMRT tumoroid models 78T and 103T after 48 h of exposure to 5 nM GTX-196 or DMSO vehicle control. **E)** UMAP analysis of metabolic gene expression profiles (n = 2,095 metabolic genes) across all primary patient samples (*n = 215*), illustrating clustering patterns by tumor type (indicated by color) and tumor subtype within a given tumor type (indicated by shape). The dataset includes AT/RT (*n = 8*; pink), ecMRT (*n = 11*; purple), SCCOHT (*n = 3*; yellow), medulloblastomas (MB; *n = 39*; green), pediatric diffuse high-grade gliomas (pHGG; *n = 39*; blue), rhabdomyosarcomas (RMS; *n = 55*; brown), and Wilms tumor samples (*n = 60*; orange). **F)** Z-score-based heatmap visualizing the average expression of genes involved in *de novo* nucleotide biosynthesis for each tumor subtype. **G)** Dose-response curves of GTX-196 for the indicated normal kidney organoid and AT/RT and SCCOHT tumoroid cultures. Data points are represented as the mean ± SD of three independent experiments, each consisting of quadruplicate measurements. Data are normalized to DMSO vehicle (100%). The grey dotted horizontal line represents a viability of 50% (IC_50_). **H)** Representative flow cytometry histograms depicting Annexin V surface expression as a marker of apoptosis in normal kidney organoids and AT/RT and SCCOHT tumoroid cultures following treatment with 50 nM GTX-196 or DMSO vehicle control. (**, *p* < 0.01; ***, *p* < 0.001; ***, *p* < 0.0001).

To do so, we performed drug screening sensitivity assays with the panel of DHODHi across multiple ecMRT tumoroid models, using normal kidney organoids as a non-malignant reference. Cellular viability was quantified using ATP levels as a surrogate for cell viability. All tested DHODHi markedly reduced ecMRT cell viability (**Figure 1B**) while having minimal impact on normal kidney organoids (**Figures S1A-E**), indicating a potentially favorable therapeutic window. The efficacy and potency of the different DHODHi varied, as reflected by their dose-response curves and corresponding IC_50_ values (**Figure 1B** and **Figures S1A-F**). GTX-196 exhibited an efficacy comparable to BAY, with an average IC_50_ of 5.1 nM ± 1.3nM, closely matching the average IC_50_ of BAY (4.7 nM, ±1.5 nM) (**Figure S1F**). Other inhibitors showed decreasing potency, with IC_50_ values increasing for PTC299 (IC_50_: 21.3 nM ±12.2 nM), AG-636 (IC_50_: 147.7 nM ±15.3 nM) and Farudodstat (IC_50_: 800.7 ±158.6 nM) (**Figure 1B** and **Figures S1A-F**). These findings indicate that GTX-196 and BAY are, at least *in vitro*, the most potent inhibitors of ecMRT growth.

Annexin V-based apoptosis assays demonstrated that GTX-196 did not induce any significant apoptotic responses in normal kidney organoids. In contrast, GTX-196 treatment produced a 2.6-fold increase in apoptosis in ecMRT tumoroids (**Figure 1C**), indicating a selective cytotoxic effect. Consistent with the known mechanism of action of DHODHi, treatment with GTX-196 led to a pronounced reduction in downstream metabolites (e.g., UTP). In parallel, upstream intermediates in the *de novo* pyrimidine biosynthesis pathway, such as N-carbamoyl-phosphate, showed significant accumulation, as measured by liquid chromatography – mass spectrometry (LC-MS)-based metabolomics (**Figure 1D**). Collectively, these data demonstrate that ecMRT are sensitive to a broad range of DHODH inhibitors, further exemplifying pyrimidine biosynthesis as a promising therapeutic candidate for ecMRT.

### Inhibition of *de novo* pyrimidine synthesis marks a vulnerability of other SWI/SNF-mutant tumors

To evaluate the broader applicability of DHODHi in pediatric cancers, we evaluated the metabolic gene expression across a range of pediatric solid (ecMRT, Wilms tumor, rhabdomyosarcoma (RMS), small cell carcinoma of the ovary, hypercalcemic type (SCCOHT)) and brain tumors (AT/RT, medulloblastoma (MB), pediatric high-grade glioma (pHGG)). Analysis of *n* = 2,095 metabolic genes in a previously published bulk RNA-seq dataset of 215 pediatric tumor samples^36^ revealed distinct clustering of brain and solid tumors in the Uniform Manifold Approximation and Projection (UMAP) plot, reflecting site-specific differences in metabolic gene expression profiles (**Figure 1E**). AT/RTs are classified into three primary molecular subtypes based on distinct DNA methylation and gene expression profiles: sonic hedgehog (AT/RT-SHH), MYC (AT/RT-MYC), and tyrosinase (AT/RT-TYR)^38^. Interestingly, two of these subtypes, AT/RT-TYR and AT/RT-MYC, did not conform to site-specific clustering and instead grouped more closely with ecMRT and SCCOHT samples, suggesting shared metabolic features across these SWI/SNF-deficient subgroups, at least on the gene expression level. Beyond this broad separation, samples also clustered according to tumor subtype within each major entity (**Figure 1E**), indicating that metabolic signatures are influenced not only by tumor location but also by underlying molecular heterogeneity.

To investigate whether SWI/SNF-mutated tumors share common metabolic features related to nucleotide metabolism, we examined the average expression of genes involved in *de novo* nucleotide biosynthesis across subtypes of various pediatric tumor types. This analysis revealed strong tumor-type-specific clustering, with SCCOHT, ecMRT, AT/RT-MYC, and AT/RT-TYR exhibiting the highest expression levels of nucleotide synthesis genes (**Figure 1F**). RMS samples displayed intermediate expression, while the expression in Wilms tumors, MB and pHGG samples was lowest. Exceptions were the blastemal subtype of Wilms tumor and the SHH subtype of MB that clustered more closely to the SWI/SNF-deficient tumors (**Figure 1F**).

Given that elevated expression of *de novo* nucleotide biosynthesis genes emerged as a common feature among SWI/SNF-mutant tumors, we next assessed the efficacy of the most potent DHODHi, BAY and GTX-196 in patient-derived AT/RT and SCCOHT tumoroids. Consistent with previous findings in ecMRT and AT/RT^22^, SCCOHT tumoroids displayed marked sensitivity to treatment with GTX-196 and BAY, with IC_50_ values in the low nanomolar range (**Figure 1G** and **Figure S1G**). This was associated with a significant increase in apoptotic cell death compared to normal kidney controls (**Figure 1H** and **Figure S1H**). Together, these results demonstrate that at least some SWI/SNF-deficient tumors share a metabolic dependency on *de novo* nucleotide synthesis, and that targeting of DHODH may represent a tissue-agnostic therapeutic vulnerability in pediatric cancers harboring inactivating alterations in SWI/SNF complex members *SMARCB1* or *SMARCA4*.

### Physiological nutrient levels attenuate the therapeutic efficacy of DHODH inhibitors

Recent studies have shown that culturing cancer cells in physiological media better reflects the metabolic state of tumors^39,40^ and improves the predictive value of *in vitro* drug responses^41,42^. However, much of our current understanding of cancer metabolism is still based on cells grown in standard cell culture media. These are not only nutrient-rich but also lack components such nucleosides and nucleobases that can potentially influence DHODHi response. To better approximate *in vivo* conditions, and thus the potential of clinically translating our findings, we evaluated the efficacy of DHODH inhibitors in cells cultured in Plasmax™, a medium designed to mimic the nutrient composition of human plasma, which contains both nucleosides and nucleobases (**Figure 2SA**)^39^. Although all tested SWI/SNF-mutant tumors were sensitive to DHODHi, we here used ecMRT tumoroid models as a proof-of-concept to evaluate physiological drug responses.

Strikingly, even at BAY and GTX-196 concentrations of up to 1 μM (∼200-fold above the IC_50_ in standard kidney organoid medium (KOM)), ecMRT tumoroids remained 20–50% viable in Plasmax™, whereas viability dropped below 10% in standard KOM (**Figures 2A,B**). Looking at both the Plasmax medium composition (**Figure S2A**) and the known mechanism of action of DHODHi, we hypothesized that the presence of uridine and uracil in Plasmax™ contributed to the reduced efficacy of BAY and GTX-196. Salvage of uridine and uracil enables cells to bypass *de novo* pyrimidine synthesis regulated by DHODH (**Figure 2C**), as cells can generate UMP via uridine-cytidine kinase (UCK), uridine phosphorylase (UPP), and/ or uracil phosphoribosyltransferase (UPRT), independent of DHODH (**Figure 2C**).

**Figure 2.**
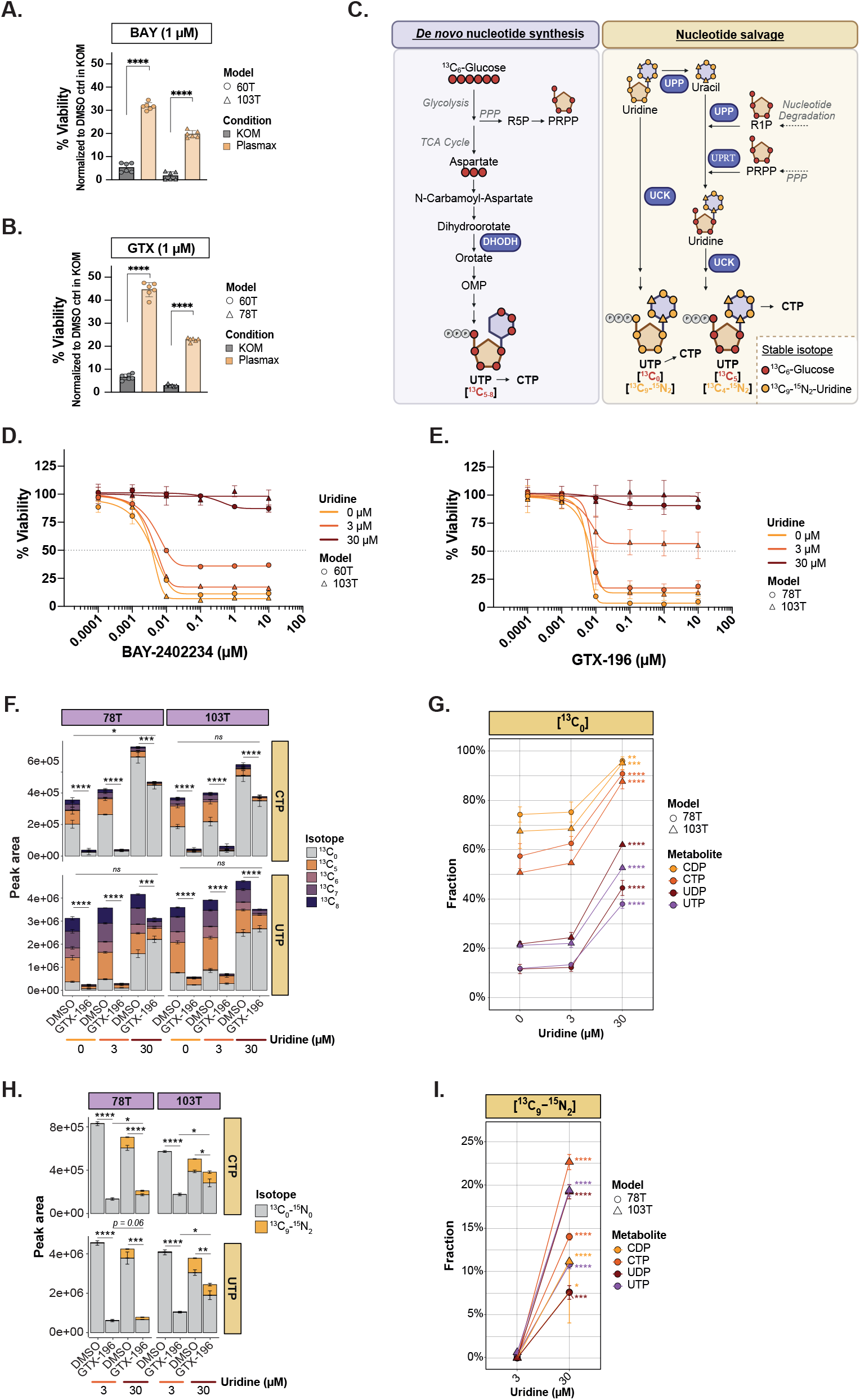
Physiological nutrient levels attenuate the therapeutic efficacy of DHODH inhibitors in ecMRT. **A-B)** Bar graphs depicting the average cell viability (%) of ecMRT tumoroid models following 120-hour treatment with 1 μM BAY-2402234 (BAY) (panel **A**) or 1 μM GTX-196 (panel **B**) in the presence of standard KOM or Plasmax™ medium. Data represent the mean ± standard deviation (SD) of n = 3 independent experiments, each performed in technical duplicates. Viability values were normalized to the DMSO vehicle control in KOM (set to 100%). **C)** Schematic overview illustrating the incorporation of carbon atoms from uniformly labeled [U-^13^C_6_]-glucose (red dots) and [U-^13^C_9_]-uridine (yellow dots) into pyrimidine nucleotide UTP via the de novo nucleotide biosynthesis and nucleotide salvage pathways. In the case of [U-^13^C_6_]-glucose, the ribose moiety (in orange) is synthesized via the pentose phosphate pathway (PPP), resulting in UTP labeled at five carbons ([^13^C_5_]). Additional carbons are contributed by aspartate (ASP) to the pyrimidine ring (in purple), generating UTP isotopologues with up to [^13^C_8_] labeling. In contrast, [^13^C_9_-^15^N_2_]-uridine is taken up through the salvage pathway as an intact molecule, yielding fully labeled UTP ([^13^C_9_-^15^N_2_]). Partial catabolism of [^13^C_9_-^15^N_2_]-uridine can lead to differential labeling patterns: UTP [^13^C_5_-^15^N_0_] indicates salvage of the labeled ribose with replacement of the pyrimidine nucleobase via de novo synthesis, while UMP [^13^C_4_-^15^N_2_] reflects incorporation of a labeled uracil base with an unlabeled ribose, suggesting base salvage following uridine breakdown. **D-E)** Dose-response curves for ecMRT tumoroid models treated with BAY (panel **D**) or GTX-196 (panel **E**) for 120 hours in standard KOM, or KOM supplemented with 3 μM uridine or 30 μM uridine. Data represent mean ± SD from three independent experiments, each consisting of technical quadruplicates. Data are normalized to DMSO vehicle in KOM (100%). The grey dotted horizontal line represents a viability of 50% (IC_50_). **F)** Total abundance and isotopic labeling pattern of CTP and UTP in two ecMRT tumoroid models following 24-hour incubation with [U-^13^C_6_]-glucose in the presence of 0⍰μM, 3⍰μM, or 30⍰μM uridine upon 48-hour treatment with DMSO vehicle or 5 nM GTX-196. Unless specified otherwise, statistical comparisons were performed within each uridine condition. **G)** Fraction of unlabeled ([^13^C_0_]) CDP, CTP, UDP, and UTP in two different ecMRT tumoroid models following 24-hour incubation with [U-^13^C_6_]-glucose in the presence of 0⍰μM, 3⍰μM, or 30⍰μM uridine. All conditions were statistically compared to the 0⍰μM uridine condition. **H)** Total abundance and isotopic labeling pattern of CTP and UTP in two ecMRT tumoroid models following 24-hour incubation with 3⍰μM or 30⍰μM [^13^C_9_-^15^N_2_]-uridine upon 48-hour treatment with DMSO vehicle or 5 nM GTX-196. Unless specified otherwise, statistical comparisons were performed within each uridine condition. **I)** Fraction of [^13^C_9_-^15^N_2_]-labelled CDP, CTP, UDP, and UTP in two different ecMRT tumoroid models following 24-hour incubation with 3⍰μM or 30⍰μM [ ^13^C_9_-^15^N_2_]-uridine. (*, *p* < 0.05; **, *p* < 0.01; ***, *p* < 0.001; ****, *p* < 0.0001).

Since uridine is a direct precursor of pyrimidine nucleotides and more readily used by cells compared to uracil^13,43^, we further delineated the contribution of uridine specifically. Hence, we treated ecMRT tumoroids with BAY and GTX-196 in standard KOM supplemented with either physiological (3 μM) or supraphysiological (30 μM) concentrations of uridine. The resulting dose-response curves in the presence of 3 μM uridine closely mirrored those observed in Plasmax™ (**Figure S2B**), suggesting that uridine contributes to the observed resistance to DHODHi in Plasmax medium. Moreover, the addition of 30 μM uridine led to near-complete rescue of cell viability following DHODH inhibition (**Figures 2D,E**). These findings indicate that extracellular uridine availability can significantly mitigate the (cytotoxic) effects of DHODHi, likely through salvage pathway engagement.

To further evaluate how uridine availability affects the balance between *de novo* nucleotide synthesis and nucleotide salvage pathway activity, we performed [U-^13^C_6_]-glucose stable isotope tracing in ecMRT tumoroids cultured in standard KOM or KOM supplemented with either physiologic (3 μM) or supraphysiologic (30 μM) uridine levels, in the presence or absence of GTX-196. Glucose-derived carbons can contribute to pyrimidine nucleotide synthesis through both the *de novo* synthesis and salvage pathways (**Figure 2C**). For either pathway, ^13^C-labelled glucose generates ribose-5-phosphate via the pentose phosphate pathway (PPP), which forms the sugar moiety of nucleotides and contributes five labeled carbons to UMP ([^13^C_5_]). Additionally, glucose-derived carbons entering the TCA cycle generate labelled aspartate ([^13^C_3_]), a key precursor for the pyrimidine ring in the *de novo* synthesis pathway. As a result, [^13^C_6-8_] isotopologues of UMP are strong indicators of active *de novo* pyrimidine synthesis. In contrast, exclusive [^13^C_5_]-labeling suggests a labelled ribose moiety was attached to a pyrimidine base derived from unlabeled sources, such as salvaged nucleobases or *de novo* synthesis from unlabeled carbon pools. Finally, [^13^C_0_] isotopologues reflect exclusive use of unlabeled precursors, consistent with uptake from pre-existing intracellular pools or salvage of extracellular uridine (**Figure 2C**). Thus, detection of high levels of [^13^C_6-8_] supports active *de novo* synthesis, whereas [^13^C_0_] is more consistent with salvage pathway utilization.

As previously described by us^22^, ecMRT cells predominantly use the *de novo* pathway for pyrimidine synthesis in the absence of exogenous uridine, as indicated by the high [^13^C_6-8_] enrichment in CTP and UTP (Figure 2F, DMSO, 0 μM uridine). Supplementation with 3 μM uridine did not markedly alter isotopic labeling patterns, whereas exposure to 30 μM uridine led to a marked increase in the unlabeled ([^13^C_0_]) fraction of pyrimidine nucleotides (**Figures 2F**, DMSO, 0-30 μM uridine; **Figure 2G**), consistent with an increased usage of the salvage pathway by ecMRT at elevated uridine concentrations. Notably, uridine supplementation did not affect the labeling of purines (**Figure S2C**), supporting the specificity of this effect for the uridine salvage route (**Figure 2C**). In addition to the altered labeling patterns, the total abundance of pyrimidine nucleotides increased under supraphysiological uridine conditions (**Figure 2F**, DMSO), suggesting enhanced pyrimidine nucleotide availability or turnover when excess uridine is present.

Treatment with GTX-196 significantly reduced pyrimidine nucleotides levels across all uridine conditions compared with DMSO controls (Figure 2F). Pyrimidine levels were almost depleted at 0 μM and 3 μM uridine after GTX-196 treatment. Yet, at 30 μM uridine, total pyrimidine levels following GTX-196 treatment remained comparable to those under standard culture conditions (DMSO, 0 μM uridine) (**Figure 2F**). Isotopic labeling indicated a shift toward salvage pathway utilization in the presence of GTX-196 at supraphysiological uridine concentrations (i.e., decreased [^13^C_6-8_], increased [^13^C_0_], Figure 2F). This shift in pathway utilization likely helps preserve total pyrimidine pools following GTX-196 treatment under high uridine conditions, which may contribute to the relative resistance of ecMRT cells to DHODH inhibition.

To directly assess uridine salvage activity and its contribution to nucleotide biosynthesis, stable isotope tracing was performed using [^13^C_9_ -^15^N_2_]-labeled uridine in ecMRT tumoroids, in the absence and presence of GTX-196. [^13^C_9_ -^15^N_2_]-uridine can give rise to multiple isotopologues in pyrimidine nucleotides, with the full [^13^C_9_ -^15^N_2_] isotopologue reflecting the incorporation of the full uridine molecule ([^13^C_5_ -^15^N_0_]-labeled ribose and [^13^C_4_ -^15^N_2_]-labeled uracil base) (**Figure 2C**). The [^13^C_4_ -^15^N_2_] isotopologue indicates incorporation of the uracil base only, without the labeled ribose. Conversely, the [^13^C_5_ -^15^N_0_] isotopologue represents incorporation of the labeled ribose alone, with an unlabeled base (**Figure 2C**).

In line with the results obtained from [U-^13^C_6_]-glucose tracing, incubation with 3⍰μM [^13^C_9_ -^15^N_2_]-uridine did not result in a measurable incorporation into pyrimidine nucleotides (**Figures 2H**, DMSO; **Figure 2I**). In contrast, 30⍰μM [^13^C_9_ -^15^N_2_]-uridine resulted in a marked increase in [^13^C_9_ -^15^N_2_]-labeledm UDP, UTP, CDP, and CTP (**Figures 2H**, DMSO; and **Figure 2I**), indicating efficient incorporation of intact exogenous uridine via the salvage pathway. At this higher uridine concentration, total pyrimidine levels were slightly lower compared with the 3⍰μM condition (**Figure 2H**, DMSO; **Figure S2D**). Although this appears to differ from the trend observed in the [U-^13^C_6_]-glucose tracing experiments, the variation likely reflects experimental context-dependent differences, as well as the inherent limitation of endpoint metabolite measurements, which capture only a snapshot of dynamic metabolic fluxes.

Treatment with the DHODHi GTX-196 led to a marked reduction in total pyrimidine nucleotide levels across all uridine conditions, although the extent of depletion was less pronounced at 30 μM uridine (**Figure 2H**), consistent with previous observations (**Figure 2F**). The comparatively higher total pyrimidine levels observed at 30 μM uridine relative to 3 μM following GTX-196 treatment (**Figure 2H**) suggest partial metabolic compensation, which may contribute to the reduced sensitivity to DHODH inhibition under uridine-rich conditions. Furthermore, at 30 μM uridine, the proportion of [^13^C_9_ -^15^N_2_]-labelled pyrimidines increased further upon GTX-196 treatment compared with DMSO controls (**Figure 2H** and **Figure S2E**), consistent with enhanced reliance on the uridine salvage pathway when *de novo* pyrimidine synthesis is impaired. Taken together, these findings indicate that ecMRT cells can utilize exogenous uridine to sustain pyrimidine pools, supporting the idea that increased salvage pathway activity contributes to DHODHi tolerance in ecMRT under uridine-rich conditions in the TME.

### ecMRT-derived interstitial fluid exhibits a metabolic composition distinct from plasma and is enriched in nucleosides and nucleobases

One limitation of using physiological plasma nutrient levels in experimental models is that tumor cells are not directly exposed to circulating plasma, but rather to the composition of extracellular fluid within the tumor microenvironment (TME), known as tumor interstitial fluid (TIF)^44^. Previous studies have reported elevated uridine levels in certain tumor tissues compared to adjacent normal tissues and plasma^45,46^, raising the question of whether a similar pattern exists in ecMRT. Besides uridine, other nucleosides and nucleobases may be present in the TME that could influence the response to DHODHi as well.

To address this, we performed metabolomic profiling on both plasma and TIF from ecMRT orthotopic xenograft models during different stages of tumor progression using LC-MS (**Figure 3A**). Principal component analyses (PCA) revealed substantial differences in the overall metabolite composition between TIF and plasma (**Figure 3B**). Moreover, hierarchical clustering of the metabolites detected across plasma and TIF samples again showed a clear separation of plasma and TIF samples, with plasma samples additionally showing a tendency to cluster according to tumor metastatic status (**Figure 3C**). Notably, many metabolites were enriched in TIF (**Figure 3C**), indicating that the ecMRT microenvironment is not uniformly nutrient-deprived, but instead contains a distinct and complex metabolic landscape that may modulate therapeutic sensitivity to DHODHi.

**Figure 3.**
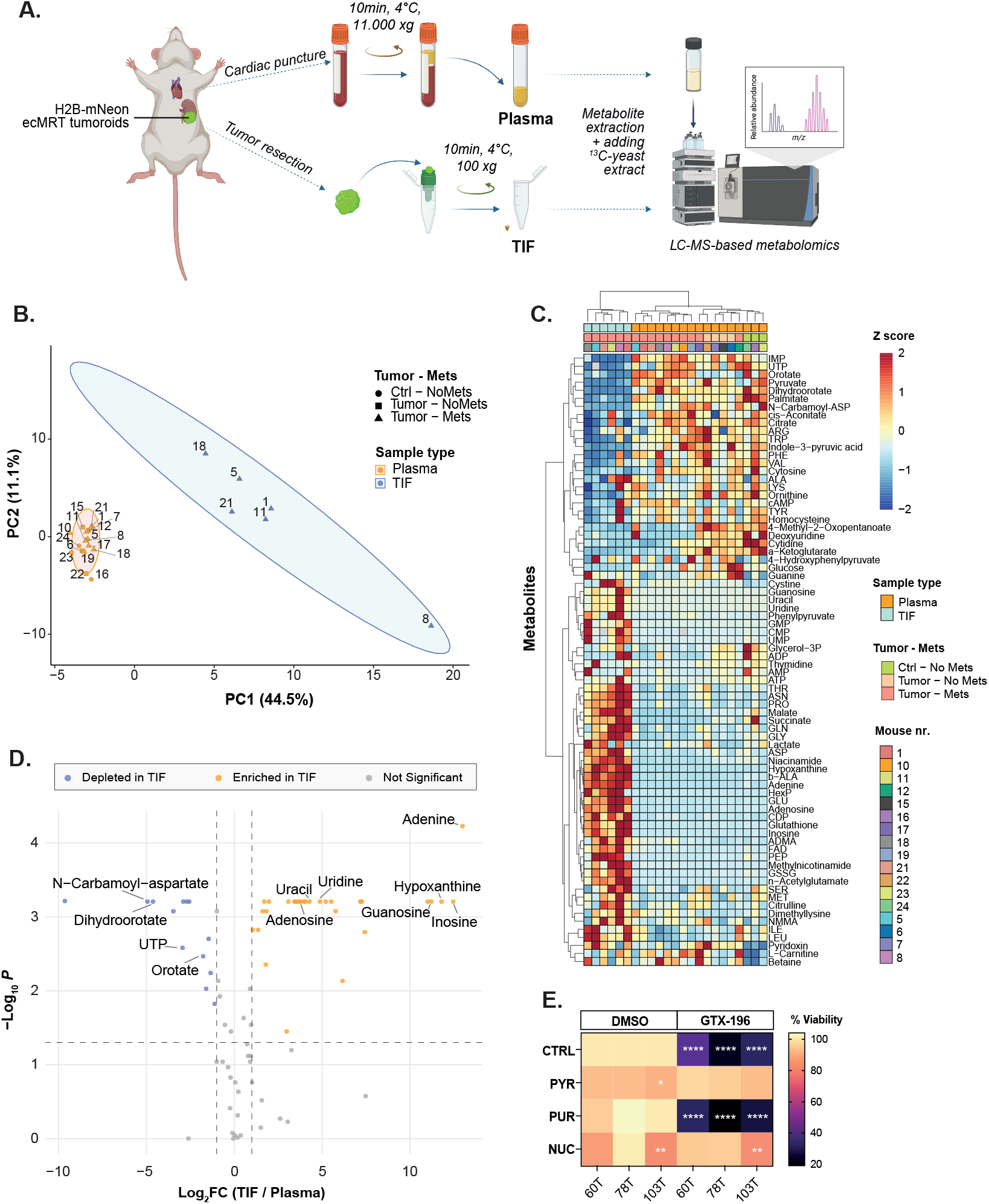
ecMRT-derived interstitial fluid exhibits a different metabolic composition and is enriched in nucleosides and nucleobases compared to plasma. **A)** Schematic representation of the workflow for isolating plasma and tumor interstitial fluid (TIF) from orthotopically transplanted ecMRT-bearing mice for LC-MS-based metabolomic profiling. **B)** Principal component analysis (PCA) of murine tumor interstitial fluid (TIF; *n = 6*, blue) and plasma samples (*n = 17*, orange), based on the abundance of *n = 76* C-yeast extract–normalized metabolites. Sample shapes indicate tumor and metastatic status: control mice without tumors (*n = 3*) are represented as circles, mice bearing only a primary tumor (*n = 4*) as squares, and mice with both primary tumors and metastases (*n = 10*) as triangles. **C)** Z score-normalized heatmap with hierarchical clustering of murine tumor interstitial fluid (TIF) and plasma samples based on the abundance of *n = 76* C-yeast extract– normalized metabolites. Annotation colors indicate sample type, tumor and metastatic status, and individual mouse identifiers. **D)** Volcano plot illustrating the log_2_ fold-change (log_2_FC) in metabolite levels between TIF and plasma samples from ecMRT-bearing mice. Control mice without tumors were excluded from this analysis. Significantly altered metabolites were defined by a log_2_FC threshold of +1 or -1 and a *p* value < 0.05. Metabolites significantly enriched in TIF are shown in orange, while those depleted in TIF are shown in blue. Selected metabolites of interest are annotated. **E)** Heatmap showing the average viability (%) of three replicates in three different ecMRT models (60T, 78T, 103T) upon treatment with 5 nM GTX-196 or DMSO vehicle control in the presence or absence of purine nucleosides (PUR; 30 μM adenosine, 30 μM guanosine), pyrimidine nucleosides (PYR; 30 μM cytidine, 30 μM uridine, 10 μM thymidine), or a mixture of both (NUC) for 120 h. Data are normalized to DMSO vehicle without nucleoside supplements (100%). (*, p < 0.05; **, p < 0.01;****, p < 0.0001).

To examine in more depth which metabolites were significantly enriched or depleted in TIF relative to plasma, we calculated the magnitude of change (log_2_ fold-change) in metabolite abundance and the corresponding statistical significance (**Figure 3D**). Key precursors involved in *de novo* pyrimidine synthesis (i.e., N-carbamoyl-aspartate, orotate, dihydroorotate) were significantly depleted in ecMRT-derived TIF compared to plasma (**Figure 3D**). In contrast, salvage pathway substrates, like purine (i.e., adenine, hypoxanthine, inosine, guanosine, adenosine) and pyrimidine (i.e., uridine, uracil) nucleosides and nucleobases were highly enriched in TIF relative to plasma (**Figure 3D**). To better characterize the amount of nucleosides and nucleobases within the TME, we compared their levels in plasma and TIF to those present in Plasmax™ (**Figure S3A**). Both uridine and uracil were markedly elevated in the biological samples relative to Plasmax™ containing 31µM uridine and 21µM uracil, whereas hypoxanthine levels were substantially reduced compared to the 51µM present in Plasmax™ (**Figure S3A**).

To test whether the elevated levels of purine and pyrimidine nucleosides detected in ecMRT-derived TIF could confer resistance to DHODHi, ecMRT tumoroids were treated with GTX-196 in standard KOM supplemented with either pyrimidine nucleosides (cytosine, uridine, thymidine), purine nucleosides (guanosine, adenosine), or a mixture of both, each provided at supraphysiological concentrations (10-30 μM). While supplementation with pyrimidine nucleosides as well as the nucleoside mixture effectively rescued the anti-proliferative effect of GTX-196, purine nucleosides could not rescue this effect (**Figure 3E**). Thus, although both purine and pyrimidine nucleosides and nucleobases were highly enriched in ecMRT-derived TIF, only pyrimidines could bypass the requirement for *de novo* nucleotide synthesis and attenuate the sensitivity of ecMRT to DHODH inhibition.

### Combined targeting of DHODH and nucleotide salvage exerts synergistic anti-tumor effects in ecMRT under (supra-)physiologic nutrient levels

Given that the presence of salvageable pyrimidine nucleosides and nucleobases within the ecMRT TME may attenuate the therapeutic efficacy of *de novo* pyrimidine synthesis inhibitors such as BAY and GTX-196, we hypothesized that simultaneous inhibition of both DHODH and nucleotide salvage pathways could restore sensitivity to DHODHi (**Figure 4A**). To test this, ecMRT tumoroids were co-treated with either BAY or GTX-196 and the FDA-approved drug Dipyridamole (DP, or Persantine)^47,48^ in the presence of uridine. DP is a known pharmacological inhibitor of human equilibrative nucleoside transporters 1 and 2 (hENT1/2), blocking nucleoside and nucleobase uptake (**Figure 4A**). Whereas monotherapy with DP had no noticeable effect on ecMRT viability (**Figure S4A**), co-treatment with DP effectively restored sensitivity of ecMRT tumoroids to DHODHi BAY and GTX-196 under Plasmax™ culture conditions (**Figure 4B, Figure S4B**). A similar resensitizing effect was observed in ecMRT tumoroids supplemented with 3 μM or 30 μM uridine, where DP induced a concentration-dependent enhancement of BAY and GTX-196 efficacy (**Figures 4C,D** and **Figures S4C,D**). Importantly, the DHODHi/DP-combo demonstrated strong synergistic activity in ecMRT tumoroids (**Figures 4E,F** and **Figures SE,F**). This synergistic interaction was further validated using two additional investigational inhibitors of hENT, draflazine and nitrobenzylthioinosine (NBMPR), which similarly enhanced the efficacy of DHODH inhibition in the presence of supraphysiological uridine levels (**Figures S4G-J**), confirming that the additional effect is exerted via hENT inhibition. Uridine is not the only salvageable metabolite present in the ecMRT TME. We therefore tested whether co-treatment with DP could restore the efficacy of DHODHi in the presence of a supraphysiological pyrimidine mixture, using a supraphysiological purine nucleoside mixture as a control. As expected, high concentrations of pyrimidine nucleosides completely abrogated the effects of GTX-196. In contrast, co-treatment with DP successfully resensitized ecMRT tumoroids to DHODHi (**Figure 4G**). A similar restoration of sensitivity to GTX-196 by DP was observed in the presence of the combined nucleoside mixture, whereas supplementation with purine nucleosides alone did not impact the efficacy of GTX-196 (**Figure 4G**), again indicating a specific role for pyrimidines in mediating resistance.

**Figure 4.**
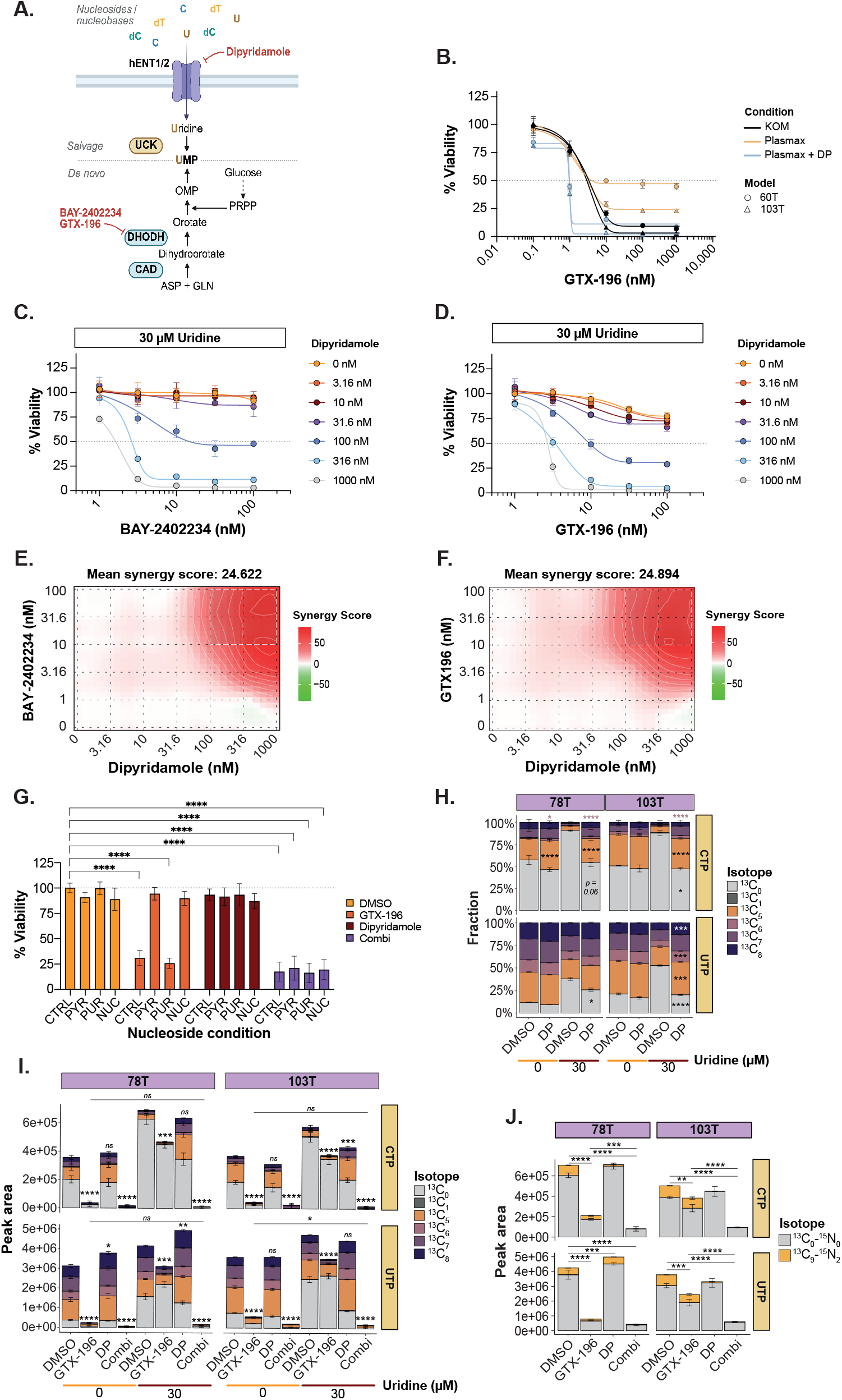
Combined targeting of DHODH and nucleotide salvage exerts synergistic anti-tumor effects in ecMRT under (supra-)physiologic nutrient levels. **A)** Schematic overview of *de novo* nucleotide biosynthesis and nucleotide salvage pathways. Nucleosides and nucleobases are imported into the cell via hENT 1 and 2 and subsequently phosphorylated to their corresponding nucleotide monophosphates (NMPs). In parallel, nucleotides can be synthesized *de novo* from metabolic precursors, resulting in the formation of NMPs. Key enzymes involved in these pathways, including uridine-cytidine kinase (UCK), DHODH, and carbamoyl-phosphate synthetase 2/ aspartate transcarbamylase/ dihydroorotase (CAD), are highlighted. Both pathways can be subjected to pharmacological inhibition by agents targeting hENT1/2 (e.g., dipyridamole) and DHODH (e.g., BAY-2402234 and GTX-196). **B)** Dose-response curves for ecMRT tumoroid models 60T and 103T treated with BAY-2402234 for 120 hours in Plasmax™ medium, with or without dipyridamole (500 nM). ecMRT tumoroids treated with BAY-2402234 in KOM medium served as a reference. Data represent mean ± SD from three independent experiments, each performed with technical duplicates. Data are normalized to DMSO vehicle in KOM (100%). The grey dotted horizontal line represents a viability of 50% (IC_50_). **C-D)** Dose-response curves for ecMRT tumoroid model 103T treated with BAY-2402234 (panel **C**) or GTX-196 (panel D) for 120 hours in KOM supplemented with 30 μM uridine, with or without different concentrations of dipyridamole. Data represent mean ± SD from three independent drug matrix experiments. Data are normalized to DMSO vehicle (100%). The grey dotted horizontal line represents a viability of 50% (IC_50_). **E-F)** ZIP^30^ synergy landscapes for BAY-2402234 + dipyridamole **(E)** and GTX-196 + dipyridamole **(F)**, corresponding to panels C–D. ZIP scores >10 indicate synergistic interactions, scores between -10 and 10 represent additive effects, and scores <10 indicate antagonism. Regions outlined in white represent the most synergistic concentration combinations. Synergy landscapes were generated using SynergyFinder software^31^. **G)** Bar graph depicting the average cell viability (%) of ecMRT tumoroid models following 120-hour treatment with 5 nM GTX-196, 10 μM dipyridamole, or their combination, in the presence or absence of nucleoside supplementation. Data represent the mean ± standard deviation (SD) of *n = 3* independent experiments, each performed in technical triplicates. Viability values were normalized to the DMSO vehicle control without nucleoside supplementation (set to 100%). **H)** Relative isotopologue distribution (fractional labeling) of CTP and UTP in two ecMRT models following 24-hour incubation with [U-^13^C_6_]-glucose upon 48-hour treatment with DMSO vehicle or 10 μM DP, in the presence or absence of 30⍰μM uridine. Statistical testing was performed within each uridine condition. I) Isotopic labeling pattern of CTP and UTP in two ecMRT tumoroid models following 24-hour incubation with [U-^13^C_6_]-glucose upon 48-hour treatment with DMSO vehicle, 5 nM GTX-196, 10 μM DP, or the combination thereof, in the presence or absence of 30⍰μM uridine. DMSO and GTX-196 reference data originate from the same [U-^13^C_6_]-glucose tracing experiment as previously presented in Figure 2, but are shown again to provide direct comparison across all treatment conditions. Unless specified otherwise, all conditions were statistically compared to the DMSO ctrl within each uridine condition. **J)** Isotopic labeling pattern of CTP and UTP in two ecMRT tumoroid models following 24-hour incubation with 30⍰μM [ ^13^C_9_-^15^N_2_]-uridine upon 48-hour treatment with DMSO vehicle, 5 nM GTX-196, 10 μM DP, or the combination thereof. DMSO and GTX-196 reference data originate from the same [ ^13^C_9_-^15^N_2_]-uridine tracing experiment as previously presented in Figure 2, but are shown again to provide direct comparison across all treatment conditions. Unless specified otherwise, all conditions were statistically compared to the DMSO ctrl. (*, *p* < 0.05; **, *p* < 0.01; ***, *p* < 0.001; ****, *p* < 0.0001).

To evaluate the impact of combining DP with DHODH inhibition on nucleotide metabolism, we performed [U-^13^C_6_]-glucose tracing in ecMRT cells treated with DMSO vehicle, GTX-196 (previously presented in **Figures 2F,G**; part of the same experimental run), DP, or the combination of GTX-196 and DP, in the presence or absence of 30 μM uridine. In line with its mechanism of action as an inhibitor of nucleoside transport, DP monotherapy did not substantially alter total pyrimidine levels or pyrimidine isotopologue distributions in the absence of exogenous uridine compared with DMSO-treated controls (**Figures 4H,I**). As shown in **Figures 2F,G**, DMSO-treated cells at 30 μM uridine displayed an increased unlabeled ([^13^C_0_]) fraction relative to the 0 μM condition, likely reflecting incorporation of exogenous uridine into pyrimidines through the salvage pathway (**Figure 4H**). Notably, treatment with DP under these high uridine conditions restored the isotopologue distribution of CTP and UTP to baseline levels (i.e., DMSO, 0 μM uridine; see **Figure 4H**), suggestive of a blocked uridine uptake and utilization. Importantly, co-treatment with DP and GTX-196 at 30 μM uridine resulted in total pyrimidine levels comparable to or lower than those observed in GTX-196-treated cells at 0 μM uridine (**Figure 4I**), indicating that inhibition of nucleoside and nucleobase transport by DP restored sensitivity to DHODH inhibition under uridine-rich conditions.

To confirm that DP treatment impairs uridine uptake and subsequent salvage into pyrimidine nucleotides in ecMRT, we performed stable isotope tracing using 30⍰μM [^13^C_9_ -^15^N_2_]-uridine following treatment with DMSO vehicle, GTX-196 (previously presented in **Figures 2H,J**; part of the same experimental run), DP, or the combination of GTX-196 and DP. In line with its proposed mechanism of action, DP treatment abrogated incorporation of the [^13^C_9_ -^15^N _2_]-label into intracellular pyrimidine nucleotide pools, while not affecting total nucleotide levels (**Figure 4J**). Moreover, measurement of extracellular uridine confirmed that [^13^C_9_ -^15^N_2_]-uridine uptake was abrogated by DP, as medium concentrations remained comparable to baseline levels in BME control samples without cells (**Figure S4K**). Notably, combined treatment with GTX-196 and DP not only completely abolished intracellular incorporation of [^13^C_9_-^15^N_2_]-label, but also significantly reduced total pyrimidine nucleotide pools (**Figure 4J**), consistent with observations from the [U-^13^C_6_]-glucose tracing experiment (**Figure 4I**). Together, these finding suggest that the DP-mediated inhibition of uridine uptake and subsequent salvage resensitizes ecMRT to DHODH inhibition.

Taken together, our findings demonstrate that a pyrimidine-enriched TME can confer resistance to DHODH inhibition, and that pharmacological inhibition of nucleoside transport effectively restores and enhances the sensitivity of ecMRT to DHODH blockade. These results provide a strong mechanistic rationale for a combinatorial therapeutic strategy targeting both *de novo* and salvage nucleotide synthesis pathways in ecMRT, a highly aggressive and currently incurable pediatric cancer.

## Discussion

To meet their elevated bioenergetic and biosynthetic demands required for sustained proliferation, tumors undergo extensive metabolic reprogramming^49^. This phenomenon, recognized as a hallmark of cancer^50^, represents a potential metabolic vulnerability that in some cases can be selectively targeted for therapeutic intervention. In our previous work^22^, we identified BAY-2402234 (BAY) as a promising therapeutic agent for malignant rhabdoid tumor (MRT), an aggressive childhood cancer with dismal prognosis. BAY is a well-characterized inhibitor of dihydroorotate dehydrogenase (DHODH), a critical enzyme in the *de novo* pyrimidine biosynthesis pathway. In the present work, we extend these findings by demonstrating that ecMRT cells also exhibit marked sensitivity to additional DHODH inhibitors (DHODHi) frequently investigated in oncologic contexts^37^, including Farudodstat (ASLAN003), PTC-299, AG636, and GTX-196. Among these, GTX-196 displayed superior potency and therapeutic efficacy in ecMRT models. Consistent with our previous observations using BAY^22^, treatment with GTX-196 robustly induced apoptosis in ecMRT cells while exerting minimal cytotoxic effects on normal kidney organoids. Collectively, these results position both BAY and GTX-196 as compelling DHODH-targeted therapeutic candidates for ecMRT, warranting further preclinical validation and clinical development.

Additionally, integrated analyses of metabolic gene expression and functional drug sensitivity assays revealed that, like ecMRT, other SWI/SNF-deficient tumors such as AT/RT and SCCOHT exhibit elevated reliance on *de novo* nucleotide biosynthesis and display a marked sensitivity to DHODHi. Similarly, other studies have highlighted a heightened sensitivity of SWI/SNF-deficient cancers to disruptions in nucleotide and, more specifically, pyrimidine pathways. For example, Metselaar *et al*.^51^ demonstrated that AT/RT cells exhibit marked sensitivity to gemcitabine, a pyrimidine nucleoside analog that interferes with DNA synthesis. Similarly, research by Young *et al*.^29^ showed that SCCOHT are vulnerable to methotrexate (MTX), a folate antagonist that impairs purine and thymidylate biosynthesis. These independent findings support a growing body of evidence that SWI/SNF-deficient malignancies harbor metabolic dependencies on nucleotide biosynthetic pathways, underscoring the therapeutic potential of targeting nucleotide metabolism in this genetically defined subset of cancers.

Although they originate from different tissues, SWI/SNF-deficient tumors share a common feature: dysregulation of the *MYC* family of oncogenes^52^, and it is therefore tempting to speculate that this common deregulation plays a role in their sensitivity to DHODHi. MYC family members (i.e., *c-Myc* (*MYC*), *N-Myc* (*MYCN*), and *L-Myc* (*MYCL*)) are functionally altered or overexpressed in the majority of human cancers^53^. In ecMRT, several studies have shown that *SMARCB1* loss drives MYC activation: enhancer rewiring increases *MYC* transcription^54^, and MYC target genes are suppressed when *SMARCB1* is reintroduced^6,55^. Similarly, AT/RT are comprised of different epigenetic subgroups, with AT/RT-SHH showing elevated *MYCN* levels, while AT/RT-MYC, similar to ecMRT, exhibits increased *MYC* expression^38^. In SCCOHTs^56^ and other *SMARCA4-*deficient cancers^57^, reintroduction of *SMARCA4* also counteracts MYC function. The tumorigenic potential of MYC stems from its broad transcriptional control over pathways that promote cellular proliferation and metabolic reprogramming, including *de novo* nucleotide biosynthesis, in part through direct regulation of DHODH^58,59^. Consistent with this, MYC-driven cancers frequently display heightened sensitivity to DHODH inhibition. Beyond ecMRT, AT/RT, and SCCOHT, other pediatric malignancies such as MYCN-amplified neuroblastoma^59,60^ and MYC-driven medulloblastoma^61^ also exhibit pronounced susceptibility to DHODHi. Conversely, DHODH inhibition has been reported to downregulate MYC expression in several tumor contexts^60–63^, suggesting reciprocal feedback between MYC activity and nucleotide metabolism that may reinforce the therapeutic vulnerability of SWI/SNF-deficient and/ or MYC-driven cancers; a phenomenon that warrants further investigation.

Our transcriptomic analyses revealed tumor-specific and subtype-specific differences in metabolic gene expression. Notably, Wilms tumors exhibited generally lower expression of nucleotide synthesis genes, whereas RMS showed higher expression; consistent with our previous BAY-2402234 drug sensitivity screening results, in which Wilms tumoroids were mainly unresponsive while RMS tumoroids showed a clear response to DHODHi^22^. Yet, the absence of healthy pediatric reference tissues limits our ability to assess whether tumors classified as “low-expressing” in our dataset truly exhibit low nucleotide biosynthetic activity. These tumors may still display elevated expression levels relative to normal tissues and therefore could remain responsive to inhibitors of nucleotide synthesis. For instance, pediatric high-grade glioma (pHGG) samples displayed comparatively lower average expression of *de novo* nucleotide biosynthesis genes in our dataset, but prior work by Pal *et al*.^64^ demonstrated that pHGGs are critically dependent on *de novo* pyrimidine synthesis, with DHODH inhibition inducing apoptotic cell death *in vitro* as well as prolonged survival of DMG-bearing mice. As such, our expression-based findings should not be interpreted as direct indicators of sensitivity or resistance to DHODH inhibition in the absence of healthy reference material and/ or functional validation.

Traditional *in vitro* culture systems often rely on non-physiologic media that fail to recapitulate the metabolic composition of the tumor microenvironment (TME), which can profoundly influence cancer cell behavior and drug sensitivity^41,42^. To elucidate the *in vivo* metabolic microenvironment of ecMRT and to more accurately recapitulate these conditions *in vitro*, we collected plasma and tumor interstitial fluid (TIF) from orthotopic ecMRT-bearing murine models and performed metabolomic profiling on these biological samples. We found that TIF is highly enriched for nucleosides and nucleobases compared to plasma. To better reflect the nutritional milieu encountered by ecMRT *in vivo*, we employed more physiologically relevant culture conditions, including defined Plasmax™ medium and suppletion of our standard medium with those nucleosides that are highly enriched in ecMRT-derived TIF. Our findings revealed that extracellular nucleosides, specifically pyrimidines, were sufficient to confer resistance to DHODH inhibition through activation of the salvage pathway; an effect that would be overlooked under standard culture conditions. The rescuing capacity of uridine, specifically in combination with DHODHi, has been previously documented by other researchers^64–67^. Importantly, this resistance could be overcome by pharmacologic inhibition of equilibrative nucleoside transporters (hENT), demonstrating how accurate modeling of the TME can uncover context-specific resistance mechanisms and identify rational combination strategies.

The metabolomic measurements of plasma and TIF represent a static snapshot of metabolite abundance within a certain tumor region, and do not capture the real-time metabolic pathway activity or spatial metabolic tumor heterogeneity. To more directly assess the functional contribution of the salvage and *de novo* nucleotide synthesis pathways *in vivo*, stable isotope tracing experiments, such as *in vivo* administration of ^13^C-labeled glucose or uridine, could be employed to map nucleotide flux in real time^68^. Additionally, including interstitial fluid from multiple tumor regions or non-malignant tissues, such as kidney interstitial fluid (KIF), would provide critical context to evaluate metabolic tumor heterogeneity as well as to more accurately distinguish tumor-specific metabolic features. Moreover, while murine models are invaluable for mechanistic studies, species-specific differences in metabolism highlight the importance of validating our findings using human ecMRT-derived TIF and plasma samples. Nevertheless, our findings underscore the importance of investigating the metabolic landscape of the TME to uncover mechanisms of drug resistance as well as to identify potential opportunities for combinatorial therapeutic strategies.

Finally, the combination of DHODH inhibition with hENT inhibition (e.g., dipyridamole) holds promising translational potential, warranting thorough evaluation of its safety, tolerability, and therapeutic window. Encouragingly, recent studies in MYCN-amplified neuroblastoma have demonstrated that this combinatorial approach is more efficacious than DHODHi or dipyridamole monotherapy in extending *in vivo* overall survival^59^. Moreover, the combination therapy was well tolerated *in vivo*, with no significant toxicity observed^59^. Building on these findings, our data provide a strong rationale for advancing preclinical and clinical investigation of this strategy in ecMRT, and potentially in AT/RT and SCCOHT, all aggressive, SWI/SNF-driven pediatric cancers for which effective treatment options remain limited.

## Methods

### Patient-derived organoids

Experiments with human material were approved by the medical ethical committee of the Erasmus Medical Center (Rotterdam, the Netherlands) and Princess Máxima Center for Pediatric Oncology (Utrecht, the Netherlands). Informed consent was obtained from the parents of all participants. Approval for use of the subject’s tissue samples within the context of this study has been granted by the Máxima biobank and data access committee (https://research.prinsesmaximacentrum.nl/en/core-facilities/biobank-clinical-research-committee) (biobank project nr. PMCLAB2018.005 and PMCLAB2018.006). Patient-derived kidney organoid cultures have been established with protocols previously described by Calandrini *et al*.^26^. Normal kidney and ecMRT tumoroids were cultured in reduced growth factor BME (Cultrex, 3533-010-02) topped with kidney organoid medium (KOM). KOM consists of AdDF+++ (Advanced DMEM/F12 containing 1x GlutaMAX, 10mM HEPES, and antibiotics; Gibco), supplemented with 1.5% B27 supplement (Gibco), 10% R-spondin-conditioned medium, EGF (50 ng/ml, Peprotech), FGF-10 (100 ng/ml, Peprotech), N-acetylcysteine (1.25 mM, Sigma), Rho-kinase inhibitor Y-27632 (10 μM, AbMole), and A83-01 (5 μM, Tocris Bioscience)^27^. KOM was changed every 3-4 days, and organoids were passaged every 1-3 weeks. Organoids were either passaged by mechanical dissociation (MRT organoids), or with TrypLE Express (Invitrogen, 1260510) with 10 μM Rho-kinase inhibitor Y-27632 (normal kidney organoids). After adding 5-10 mL AdDF+++ and centrifugation at 300 rcf, cells were reseeded in BME and topped with KOM. Atypical teratoid/rhabdoid tumoroid and small cell carcinoma of the ovary, hypercalcemic type (SCCOHT) tumoroid cultures were maintained as described by Paassen *et al*.^28^ and Kim *et al*.^29^, respectively.

### Drug screens

Drug screening assays on AT/RT and SCCOHT tumoroids were performed according to the methodology outlined by Paassen *et al*.^28^ and Kim *et al*.^29^. Drug screens on malignant rhabdoid tumor (MRT) tumoroids and normal kidney organoids were carried out following the protocol described by Kes *et al*.^22^.

Briefly, between 500 (for normal kidney and ecMRT organoids) and 2.000 (for AT/RT and SCCOHT tumoroids) small organoids were seeded per well in black 384-well plates (Corning) in a total volume of 40 μL. Normal kidney and ecMRT organoids were embedded in a 5% BME slurry in kidney organoid medium (KOM), while AT/RT and SCCOHT tumoroids were seeded in suspension in their respective tumor-specific media: TSM (tumor stem medium) for AT/RT and SCCOHT medium for SCCOHT. For uridine rescue experiments, 3 μM or 30 μM uridine (Sigma-Aldrich, U3750) was added to the KOM. Seeding of the organoids was done with a Multi-drop Combi Reagent 8 Dispenser (Thermo Scientific). Compound treatments were applied using a Tecan D300e Digital Dispenser. BAY-2402234 (MedChemExpress, HY-112645), GTX-196 (Genase Therapeutics), AG-636 (MedChemExpress, HY-137463), ASLAN003/Farudodstat (MedChemExpress, HY-129239), PTC299 (MedChemExpress, HY-124593), Dipyridamole (MedChemExpress, HY-B0312), Nitrobenzylthioinosine (NBMPR; MedChemExpress, HY-W010936), and Draflazine (MedChemExpress, HY-106841) were tested at final concentrations ranging between 0.01 nM and 10 μM. The Tecan D300e dispenser was used to normalize drug concentrations for dimethyl sulfoxide (DMSO) content. Cells treated with DMSO served as negative controls. For standard dose–response experiments, three biologically independent experiments were conducted for each organoid model, with four technical replicates per condition within each experiment. For drug combination (matrix) studies, three independent drug matrices were screened per organoid model. 120 hours after adding the drugs, ATP levels were assessed with CellTiter-Glo 3D reagent (Promega) according to the manufacturer’s instructions on a BMG Labtech FLUOstar® Omega microplate reader. Results were normalized to DMSO vehicle (100%). Survival data was analyzed and visualized in GraphPad Prism (version 10.4.1). IC_50_ values were calculated in RStudio, using the *drc* package (v.3.0-1). Synergy ZIP scores^30^ and plots were generated using the SynergyFinder tool^31^ (https://synergyfinder.org/).

### Flow cytometric analyses of Annexin V/DAPI positive cells

A modified version of the protocol described by Kes *et al*.^22^ was utilized for this experiment. Normal kidney organoids (JD065H, 60H, 103H), ecMRT tumoroids (60T, 78T, 103T), AT/RT tumoroids (ATRT-04, ATRT-15, ATRT-06, ATRT-08) and SCCOHT tumoroids (OVT-029, OVT-039) were passaged and plated in 75% BME droplets topped up with KOM (normal kidney organoids and ecMRT tumoroids) or in suspension (AT/RT and SCCOHT tumoroids) in their designated medium. After three days (MRT, AT/RT, and SCCOHT tumoroids) or six days (normal kidney organoids), cells were re-plated in 5% BME slurry with KOM (normal kidney organoids and ecMRT tumoroids) or in suspension (AT/RT and SCCOHT tumoroids) and treated with either DMSO vehicle or 50 nM GTX-196 (Genase Therapeutics). Following 120 hours of treatment, organoids and supernatants were harvested and dissociated into single-cell suspensions using TrypLE Express (ThermoFisher) in the presence of Rho-kinase inhibitor Y-27632 (AbMole). Single-cell suspensions were stained with APC-Annexin V (BD Biosciences, #550475) and DAPI (ThermoFisher, #D1306) in Annexin V binding buffer supplemented with 2.5 mM Ca^2+^. Flow cytometric analysis was performed using the Beckman Cytoflex LX, and data were analyzed with FlowJo software (BD Biosciences, version 10.8.2). Apoptotic indices were calculated by normalizing the percentage of apoptotic cells to DMSO controls (set to 1). Live and apoptotic fractions were determined by dividing the number of events in each gated population by the total number of measured events (Live + Early Apoptotic + Late Apoptotic = 1).

### Plasmax rescue experiments

Organoids were dissociated and made into a single-cell solution using mechanical disruption. Single cells were plated at a density of 2.000 cells/μL in 5 μL 75% BME droplets topped with KOM^PL^ and Plasmax (CancerTools, 156371) (see exact formulations in Table 1) in a pre-warmed flat-bottom 96-well plate (Greiner, 655-160). After 72 hours, treatment with 0.01 nM – 1 μM GTX-196 (Genase Therapeutics) or BAY-2402234 (MedChemExpress, HY-112645), alone or in combination with 500 nM Dipyridamole (MedChem Express, HY-B0312), was started. In all the wells, the medium was refreshed daily, due to the rapid nutrient depletion in physiological Plasmax medium. All treatments lasted for 120 hours. On the day of the readout, all medium was removed and 45 μL of adDF+++ together with 50 μL of the CellTiter-Glo® 3D solution (Promega) was added to each well. After 5min of shaking and 30 min at RT in the dark, 80 μL of each well was transferred into a black clear clear-bottom 396-well plate (Corning). Luminescence was measured with the FLUOstar® Omega microplate reader (BMG Labtech), and data were analyzed with GraphPad Prism software (version 10.4.1). Results were normalized to DMSO-only treated cells screened in KOM (100%) for each separate organoid model. Per organoid model, three independent experiments were performed, using technical duplicates for each experimental condition.

**Table 1.** Media formulations.

| Component | KOM<sup>PL</sup> | Plasmax | [$^{13}\text{C}_6$]-glucose KOM | [$^{13}\text{C}_9$-$^{15}\text{N}_2$]-uridine KOM |
| --- | --- | --- | --- | --- |
| **Base medium** | Advanced DMEM/F12 | Plasmax™ | SILAC Advanced DMEM/F12 | Advanced DMEM/F12 |
| **GlutaMAX** | 1x (1:100) | NA | NA | NA |
| **Pen/Strep** | 1x (1:100) | 1x (1:100) | 1x (1:100) | 1x (1:100) |
| **R-spondin-conditioned medium** | 10% | 10% | 10% | 10% |
| **B27** | 1.5% | 1.5% | 1.5% | 1.5% |
| **N-acetylcysteine** | 1.25 mM | 1.25 mM | 1.25 mM | 1.25 mM |
| **Rho-kinase inhibitor Y-27632** | 10 μM | 10 μM | 10 μM | 10 μM |
| **A83-01** | 5 μM | 5 μM | 5 μM | 5 μM |
| **EGF** | 50 ng/mL | 50 ng/mL | 50 ng/mL | 50 ng/mL |
| **FGF-10** | 100 ng/mL | 100 ng/mL | 100 ng/mL | 100 ng/mL |
| **D-Glucose** | NA | NA | NA | 17.5 mM |
| **[U-$^{13}\text{C}_6$]-glucose** | NA | NA | 17.5 mM | NA |
| **L-Glutamine** | NA | NA | 2 mM | 2 mM |
| **L-Arginine** | NA | NA | 0.7 mM | NA |
| **L-Lysine** | NA | NA | 0.5 mM | NA |
| **Uridine** | ± 3 μM | NA | ± 3 μM or ± 30 μM | NA |
| **[$^{15}\text{N}_2$-$^{13}\text{C}_9$]-uridine** | NA | NA | NA | ± 3 μM or ± 30 μM |

### Nucleoside rescue experiments

Organoids were dissociated and made into a single-cell solution using mechanical disruption. Single cells were plated at a density of 2.000 cells/μL in 5 μL 75% BME droplets topped with KOM in a pre-warmed flat-bottom 96-well plate (Greiner, 655-160). After 72 hours of initial culture, cells were treated with 5 nM GTX (Genase Therapeutics), 10 μM dipyridamole (MedChem Express, HY-B0312), or a combination of both. These treatments were administered either alone or in conjunction with purine nucleosides (30 μM adenosine [A4036] and 30 μM guanosine [G6264], Sigma-Aldrich), pyrimidine nucleosides (30 μM cytidine [C4654], 10 μM thymidine [T1895], and 30 μM uridine [U3750], Sigma-Aldrich), or a combined nucleoside mixture. To account for the chemical instability of nucleosides in culture conditions, the medium in all wells was refreshed daily. All treatments lasted for 120 hours. On the day of the readout, all medium was removed and 45 μL of adDF+++ together with 50 μL of the CellTiter-Glo® 3D solution (Promega) was added to each well. After 5min of shaking and 30 min at RT in the dark, 80 μL of each well was transferred into a black clear clear-bottom 396-well plate (Corning). Luminescence was measured with the FLUOstar® Omega microplate reader (BMG Labtech), and data were analyzed with GraphPad Prism software (version 10.4.1). Results were normalized to DMSO-only treated cells (100%) for each separate organoid model. Three independent experiments were performed, using technical triplicates for each experimental condition.

### Stable isotope tracing in ecMRT tumoroids

MRT tumoroids 78T and 103T were processed into single cell suspensions as described in previous sections. For each condition, 0.25×10^6^ cells were plated in triplicate in 75% BME droplets in KOM medium supplemented with or without 3 μM or 30 μM uridine (see Table 1). 72 hours after seeding, cells were treated with dimethyl sulfoxide (DMSO, vehicle control), 5 nM GTX-196 (Genase Therapeutics), 10 μM Dipyridamole (DP; MedChem Express, HY-B0312), or a combination of 5 nM GTX-196 and 10 μM DP for 48 hours. One day after the start of treatment, 24-hour isotopic labelling with [U-^13^C_6_]-glucose (Cambridge Isotopes, CLM-1396-PK) or [^15^N_2_ -^13^C_9_]-uridine was initiated. For isotopic labeling with [U-^13^C_6_]-glucose, the culture media were replaced with glucose-free SILAC Advanced DMEM/F-12 (Gibco, A2494301). These media were supplemented with 17.5 mM [U-^13^C_6_]-glucose and additional components outlined in Table 1. For isotopic labeling with [^15^N_2_ -^13^C_9_]-uridine, standard culture media – Advanced DMEM/F12 – was used and supplemented with 3 μM or 30 μM [U-^15^N_2_ -^13^C_9_]-uridine and additional components as indicated in Table 1. Since BME compositions could change after extended culturing, BME containing no cells was plated in triplicate for each tumor type to correct for possible background effects. 24 hours after labelling, 10 µL of conditioned medium was transferred to 1.5 mL Eppendorf tubes and 900 µL ice-cold MS lysis solvent (LS) composed of methanol/acetonitrile/Milli-Q (2:2:1) was added to the tubes. After that, medium was removed from the wells and all the wells were washed with 1 mL cold PBS, without disrupting the BME droplets. PBS was removed, and 500 µL ice-cold LS was added to each well. Plates were put on a plate Rocker at 4°C for 10min to induce lysis. After 10min, LS was removed from the wells and collected in 1.5 mL Eppendorf tubes that were then put in a Thermoshaker at 4°C for 15 min. Samples were cleared by centrifugation at 12.000 rcf for 15 min. Supernatants were frozen at -80°C until further analyses by LC-MS-based metabolomics.

### Animal studies

Mouse experiments were conducted in accordance with the Dutch Experiments on Animals Act 2014 and approved by the Animal Welfare Body of the Princess Máxima Center for Pediatric Oncology (AVD39900 2022 16405, PmcDro16405.03 and PmcDro16405.04). 7 to 14-week-old NOD.Cg-*Prkdc*^*scid*^*Il2rg*^*tm1Wjl*^/SzJ (NSG) mice (male/female) were used as acceptors for orthotopic transplantation. Mice were kept under standard temperature and humidity conditions in individually ventilated cages (Innovive), with food and water provided *ad libitum*. Before orthotopic xenotransplantation, patient-derived ecMRT of the kidney organoids of model 60T were lentivirally transduced to stably express luciferase (pLKO-1.UbC.fire-fly-luciferase.blast)^32^ to follow tumor progression *in vivo* using bioluminescence imaging (BLI) as well as H2B-mNeon (pLKO-1.EF1A.Puro.P2A.H2B-mNeon)^33^ to perform *ex vivo* tumor staging using fluorescence stereo microscopy (Leica, M205 FA). ecMRT organoids were implanted under the renal capsule of the left kidney in NSG mice. Tumor growth was monitored every three weeks via BLI starting 21 days post-implantation. D-luciferin (150 mg/kg) was administered subcutaneously before imaging, followed by anesthesia with isoflurane and placement in a dark chamber for bioluminescent imaging (MILabs, Optical Imaging System). An extra BLI session was performed under terminal anesthesia if the animal had to be sacrificed due to human endpoint. Blood collection was performed under isoflurane anesthesia, after which mice were euthanized by cervical dislocation, followed by the collection of blood and tissue/organs. Mice were sacrificed at different time points after orthotopic transplantation (3 weeks, 6 weeks, or at humane endpoint after 12-18 weeks) to follow tumor progression over time. Final tumor staging (no tumor, primary tumor only, or primary tumor with metastases) was determined by assessing H2B-mNeon signal in harvested tissues and organs. Information on all mice, including tumor staging and the collected samples, is provided in Table S1.

### Tumor interstitial fluid (TIF) and plasma collection for metabolite extraction

Blood of tumor-bearing mice was collected under isoflurane anesthesia, after which mice were euthanized by cervical dislocation. Tumors and other organs were rapidly excised and placed on ice. Whole blood was obtained via terminal cardiac puncture and transferred into 1.5 mL EDTA-coated microcentrifuge tubes (Eppendorf). Blood samples were centrifuged at 11.000 rcf for 10 min at 4°C to isolate plasma, which was aliquoted and stored at −80°C for subsequent analyses.

Excised tumors were sectioned into approximately 5–10 mm^2^ fragments, briefly rinsed in sterile Dulbecco’s Phosphate-Buffered Saline (DPBS; Gibco, 14080055), and gently blotted on filter paper (Whatman). Tissue fragments were then placed onto 20 µm pluriStrainer Mini Cell Strainer filters (pluriSelect, 43-10020-50) mounted atop sterile 1.5 mL microcentrifuge tubes (Eppendorf). The assembly was centrifuged at 100 rcf for 10 min at 4°C to facilitate tumor interstitial fluid (TIF) extraction.

The collected TIF was subsequently transferred to sterile 0.5 mL microcentrifuge tubes (Eppendorf) and stored at −80°C until further use. TIF volumes ranging from 5 to 50 μL were successfully isolated from *n* = 6 primary renal tumors (see details in Table S1). In the remaining cohort, primary tumor size was insufficient for fluid extraction. The yield of TIF per tumor was consistent with previously reported methodologies for murine tumor models^34^.

Polar metabolites were extracted from plasma and TIF samples using a methanol/acetonitrile/Milli-Q extraction solvent (2:2:0.5; HPLC-grade), including a ^13^C-labeled yeast extract as an internal standard (Cambridge Isotope Laboratories, Andover, MA; ISO1). Briefly, 5⍰μL of the sample was mixed with 45⍰μL of the extraction solvent in 1.5⍰mL microcentrifuge tubes (Eppendorf). The mixtures were incubated in a 4 °C thermoshaker for 10 min to facilitate metabolite extraction, followed by centrifugation at 12.000 rcf for 15 min at 4 °C to pellet insoluble material. An aliquot of 20⍰μL of the resulting supernatant containing polar metabolites was collected and used for subsequent liquid chromatography–mass spectrometry (LC-MS) metabolomics analysis.

### Liquid chromatography – mass spectrometry (LC-MS)-based metabolomics

LC-MS analyses of metabolites were performed as previously described^22,35^ on a Q Exactive HF mass spectrometer (Thermo Scientific) coupled to a Vanquish autosampler and pump (Thermo Scientific). Separation of metabolites was done using a Sequant ZIC-pHILIC column (2.1 × 150 mm, 5 μm; Merck) coupled to a ZIC-pHILIC guard column (2.1 × 20 mm, 5 μm; Merck), using elution buffers acetonitrile for A, and 20mM (NH_4_)_2_CO_3_, 0.1% NH_4_OH in LC/MS grade water (Biosolve) for B. Column temperature was 30°C. Flow rates were set at 100 μl/min and a gradient ran from 80%A to 20%A. Sample injection volumes were 5 µL. The MS operated in polarity-switching mode with spray voltages of -3.5 kV and 4.5 kV. Sheath gas, auxiliary gas, and sweep gas flow rates were 35, 10, and 1 units, respectively.

Data was analyzed using TraceFinder software (Thermo Scientific). Identification and quantification of metabolites was based on exact mass within 5 ppm and validated further by concordance with m/z, retention times, and peak shape of reference standards of *n*=132 metabolites of interest that were included in the same run.

For the stable isotope tracing experiments in ecMRT tumoroids, peak intensities were normalized based on total ion count, and distributions of isotopes were corrected for the natural abundance of ^13^C.

For metabolomic analyses of plasma and TIF, metabolite signals were normalized to the corresponding ^13^C-labelled metabolite in the internal standard (^13^C-labeled yeast extract) to correct for matrix effects. Specifically, the peak area of the (unlabeled) metabolite in the analyte was divided by the signal of the corresponding (fully ^13^C labelled) metabolite in the internal standard:

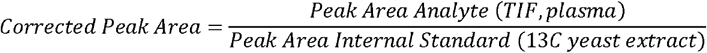

In cases where a labeled version of the exact metabolite could not be identified in the sample, a closely related labeled metabolite—referred to as a “proxy metabolite”—was used instead. This proxy was chosen based on its similar behavior during LC-MS analysis (e.g., similar retention time and/or biochemical properties). See **Table S2** for the labeled metabolites paired to each metabolite analyzed without an internal standard. *N* = 76 metabolites could be normalized via this method.

### Bulk RNA-seq analyses

For gene expression analyses, bulk RNA-seq data were obtained from the publicly available repository hosted on Zenodo related to the publication of the M&M classifier^36^. Data were subsetted to only include extracranial MRT (ecMRT), atypical teratoid rhabdoid tumors (AT/RT), Medulloblastoma (MB), pediatric high-grade glioma (pHGG), Rhabdomyosarcoma (RMS), small cell carcinoma of the ovary, hypercalcemic type (SCCOHT), and Wilms tumors (Wilms). Data processing, normalization, and visualization were performed using R (version 4.4.0). Before downstream analyses, genes with fewer than 10 read counts across all samples were excluded to reduce background noise and improve data quality. Raw count data were normalized using the variance-stabilizing transformation (VST) implemented in the *DESeq2* package to account for differences in sequencing depth and variance across samples. Dimensionality reduction was carried out using Uniform Manifold Approximation and Projection (UMAP) via the *umap* package, and results were visualized with the *ggplot2* package. Heatmaps were generated using the *pheatmap* package.

### Statistical analyses

To compare the proportions of live, early apoptotic, and late apoptotic cells between DMSO- and GTX-196–treated organoids, multiple paired Student’s *t* tests were performed. Differences in cell viability between GTX-196- and BAY-2402234-treated samples in standard KOM versus Plasmax™ medium were assessed using unpaired Student’s *t* tests. Viability differences between DMSO-treated controls (no nucleosides) and DMSO-, GTX-196-, Dipyridamole (DP)-, or combi-treated samples with or without nucleoside supplementation were evaluated using two-way ANOVA with Dunnett’s correction for multiple comparisons. Comparisons of metabolite levels between tumor interstitial fluid (TIF) and plasma from ecMRT-bearing mice were conducted using two-sided Wilcoxon rank-sum tests. For multi-group comparisons of metabolite levels based on sample type (TIF or plasma) and metastatic status, Kruskal-Wallis tests were applied. Total metabolite peak areas between DMSO- and GTX-196-treated ecMRT tumoroids were compared using two-sided unpaired Student’s *t te*sts and corrected for multiple testing with the Benjamini-Hoch correction. To compare the difference in isotopologue fractions and total metabolite peak areas for ecMRT models cultured in medium containing 3 μM versus 30 μM labelled uridine, a Student’s t test was performed, and *p* values were adjusted for multiple testing using the Benjamini-Hoch correction. Statistical significance of differences in isotopologue fractions and total metabolite peak areas across the various uridine and treatment conditions was evaluated using One-way ANOVA, followed by Tukey’s honestly significant difference (HSD) post hoc test for multiple comparisons. *P* values < 0.05 were considered significant. All statistical data can be found in the figure legends.

## Acknowledgments

We are grateful to the patients and parents who agreed to participate in our research. We would like to thank the Princess Máxima Center Biobank and Data Access Committee for providing tissues and data. We thank Allard Kaptein and Ernst Geutjes (Genase Therapeutics) for generously providing the compound GTX-196 for research purposes. C.R.B. and J.D. received funding from Foundation Children Cancer-free (KiKa #377), and J.D. from Oncode Accelerator, a Dutch National Growth Fund project under grant number NGFOP2201.

## Supplementary information

**Supplementary Figure 1.**
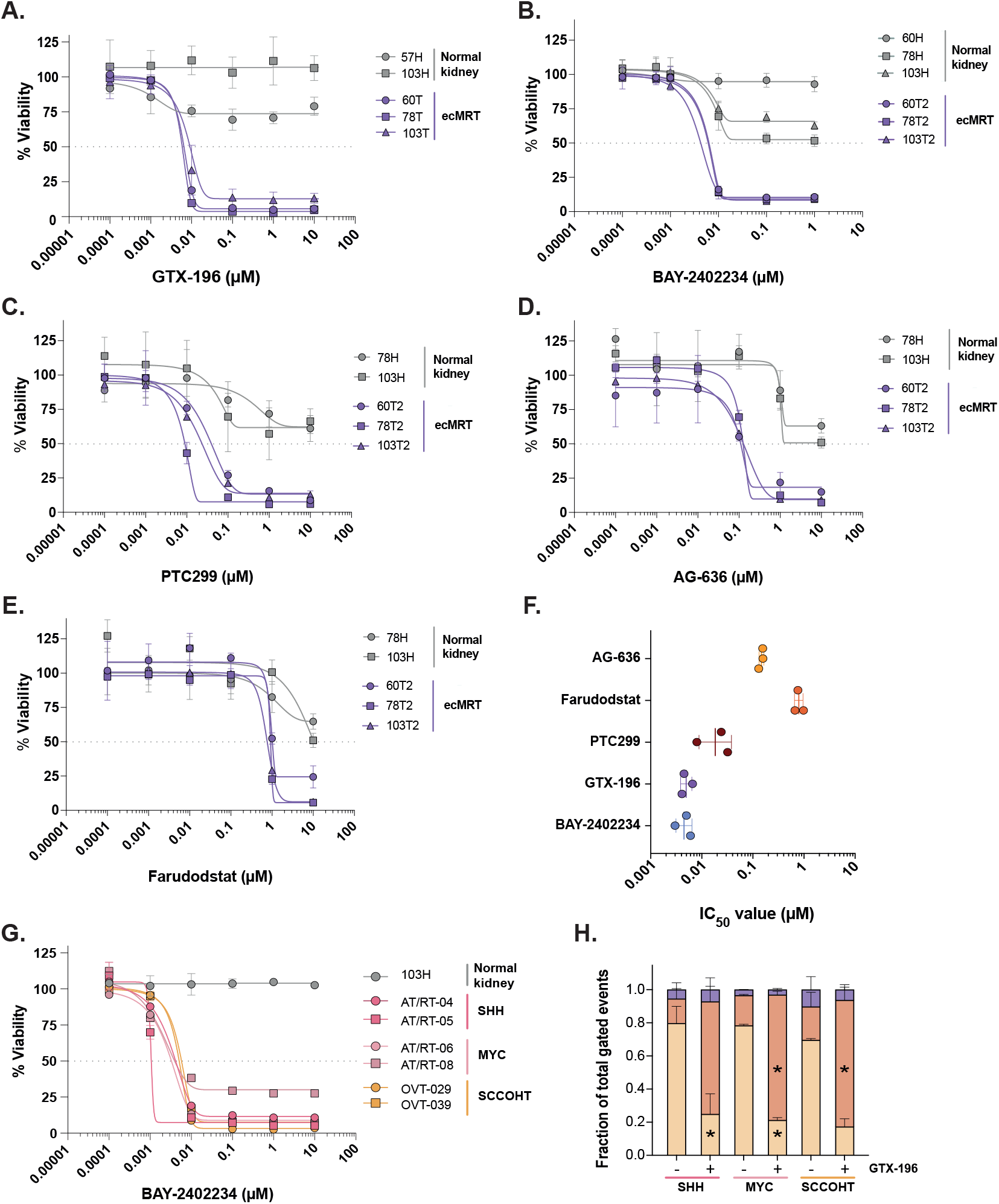
Inhibition of de novo pyrimidine synthesis enzyme DHODH marks a vulnerability of SWI/SNF-mutant tumors. **A-E)** Dose-response curves of DHODH inhibitors GTX-196 (panel **A**), BAY-2402234 (panel **B**), PTC299 (panel **C)**, AG-636 (panel **D)**, and Farudodstat (ASLAN003; panel **E)** for the indicated normal kidney organoid and ecMRT tumoroid cultures. Data points are represented as the mean ± SD of quadruplicate measurements. Data are normalized to DMSO vehicle (100%). The grey dotted horizontal line represents a viability of 50% (IC_50_). **F)** Scatterplots showing the individual IC_50_ of each ecMRT tumoroid model, grouped per DHODH inhibitor. Mean ± SD are also plotted. **G)** Dose-response curves of BAY-2402234 for the indicated normal kidney organoid and AT/RT and SCCOHT tumoroid cultures. Data points are represented as the mean ± SD of three independent experiments, each consisting of quadruplicate measurements. Data are normalized to DMSO vehicle (100%). The grey dotted horizontal line represents a viability of 50% (IC_50_). **H)** Bar graphs representing live (yellow), early apoptotic (orange), and late apoptotic (purple) cell fractions of AT/RT-SHH, AT/RT-MYC and SCCOHT tumoroids upon treatment with 50 nM GTX-196 or DMSO vehicle control for 120 h. The means ± SD of *n = 2* models are plotted. (*, *p* < 0.05).

**Supplementary Figure 2.**
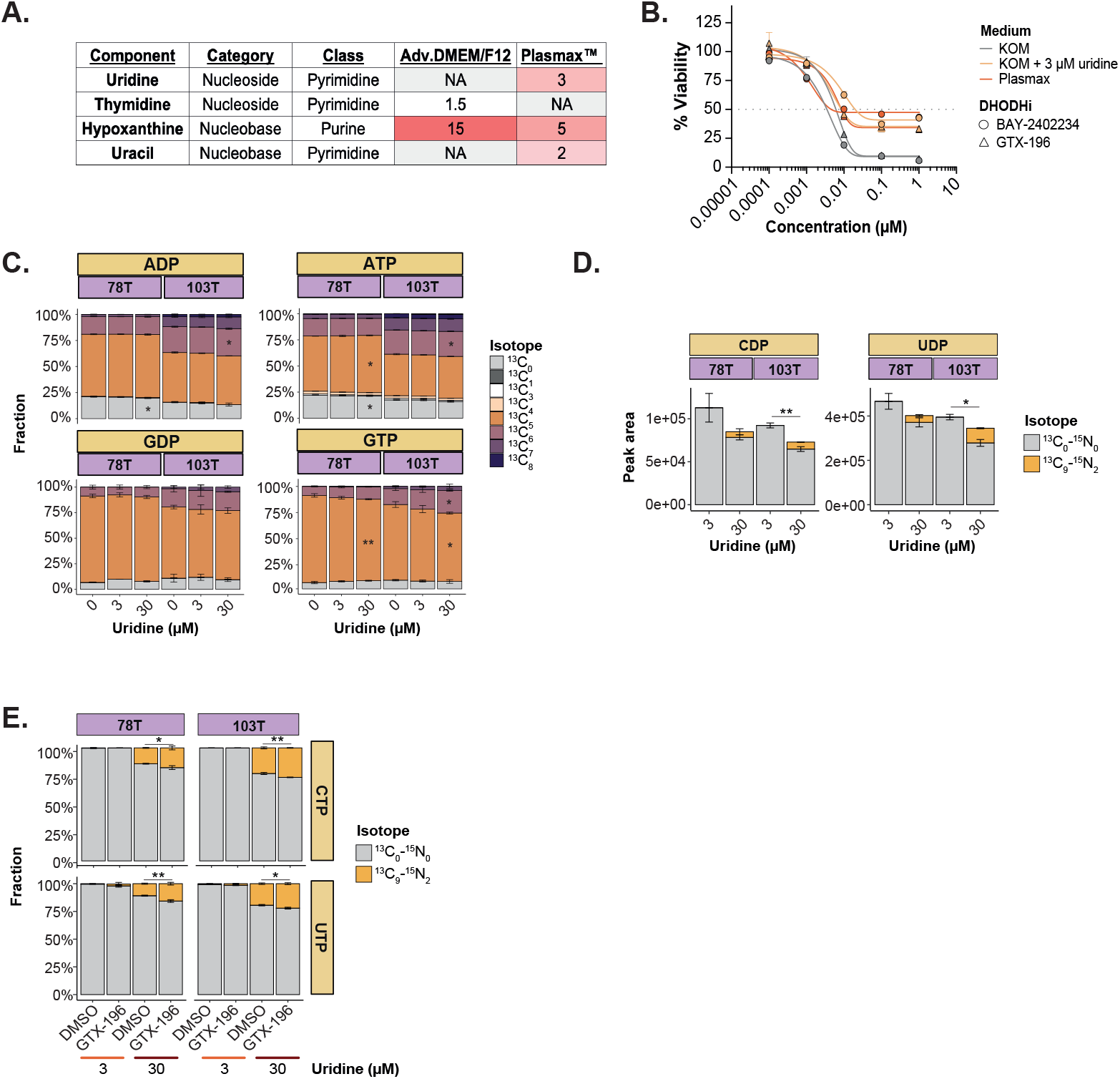
Physiological nutrient levels attenuate the therapeutic efficacy of DHODH inhibitors in ecMRT. **A)** Overview of the concentrations of nucleosides and nucleobases in Advanced DMEM/F12 (base medium for KOM) and Plasmax™ according to the manufacturer. “NA” indicates that the respective compound is absent from the medium formulation. Shades of red represent relative concentrations of each component. **B)** Dose-response curves for ecMRT tumoroid model 60T treated with BAY or GTX-196 for 120 hours in standard KOM, KOM supplemented with 3 μM uridine, or Plasmax. Data represent mean ± SD from three independent experiments. Data are normalized to DMSO vehicle in KOM (100%). The grey dotted horizontal line represents a viability of 50% (IC_50_). **C)** Relative abundance of different isotopologues in ADP, ATP, GDP, and GTP in two different ecMRT tumoroid models following 24-hour incubation with [U-^13^C_6_]-glucose in the presence of 0⍰μM, 3⍰μM, or 30⍰μM uridine. All conditions were statistically compared to the 0⍰μM uridine condition. **D)** Total abundance and isotopic labeling pattern of CDP and UDP in two ecMRT tumoroid models following 24-hour incubation with 3⍰μM or 30⍰μM [ ^13^C_9_-^15^N_2_]-uridine. Statistical testing was performed within each uridine condition. **E)** Isotopic labeling pattern of CTP and UTP in two ecMRT tumoroid models following 24-hour incubation with 3⍰μM or 30⍰μM [ ^13^C_9_-^15^N_2_]-uridine upon 48-hour treatment with 5 nM GTX-196 or DMSO vehicle. Unless specified otherwise, all conditions were statistically compared to the DMSO ctrl within each uridine condition. (*, *p* < 0.05; **, *p* < 0.01; ***, *p* < 0.001; ****, *p* < 0.0001).

**Supplementary Figure 3.**
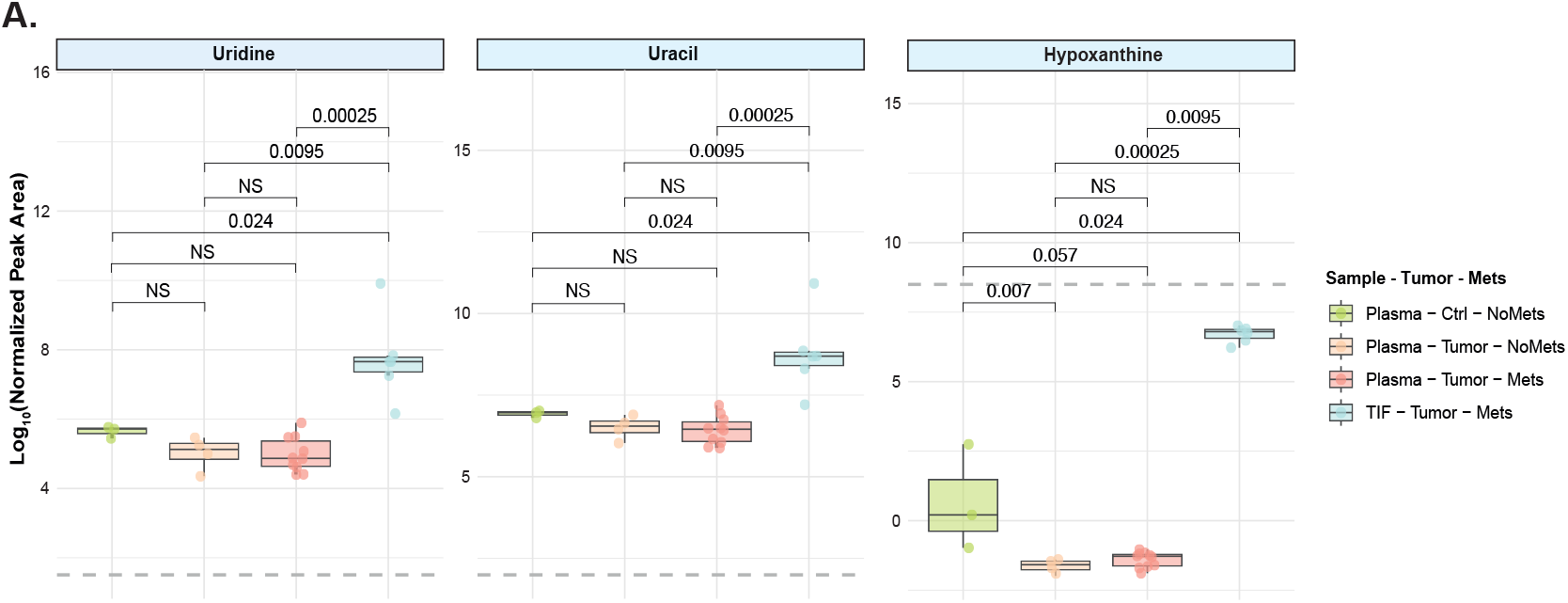
ecMRT-derived interstitial fluid exhibits a different metabolic composition and is enriched in nucleosides and nucleobases compared to plasma. **A)** Box-plots displaying the log_10_-transformed ^13^C-yeast extract-normalized peak areas of the metabolites uridine (left), uracil (middle), and hypoxanthine (right). See **Table S2** for proxy-metabolites used for normalization. Each dot represents the normalized peak area from an individual sample. Sample colors correspond to sample type and tumor metastatic status. The gray dashed line indicates the log_10_-normalized peak area measured in Plasmax™ reference samples containing known metabolite concentrations.

**Supplementary Figure 4.**
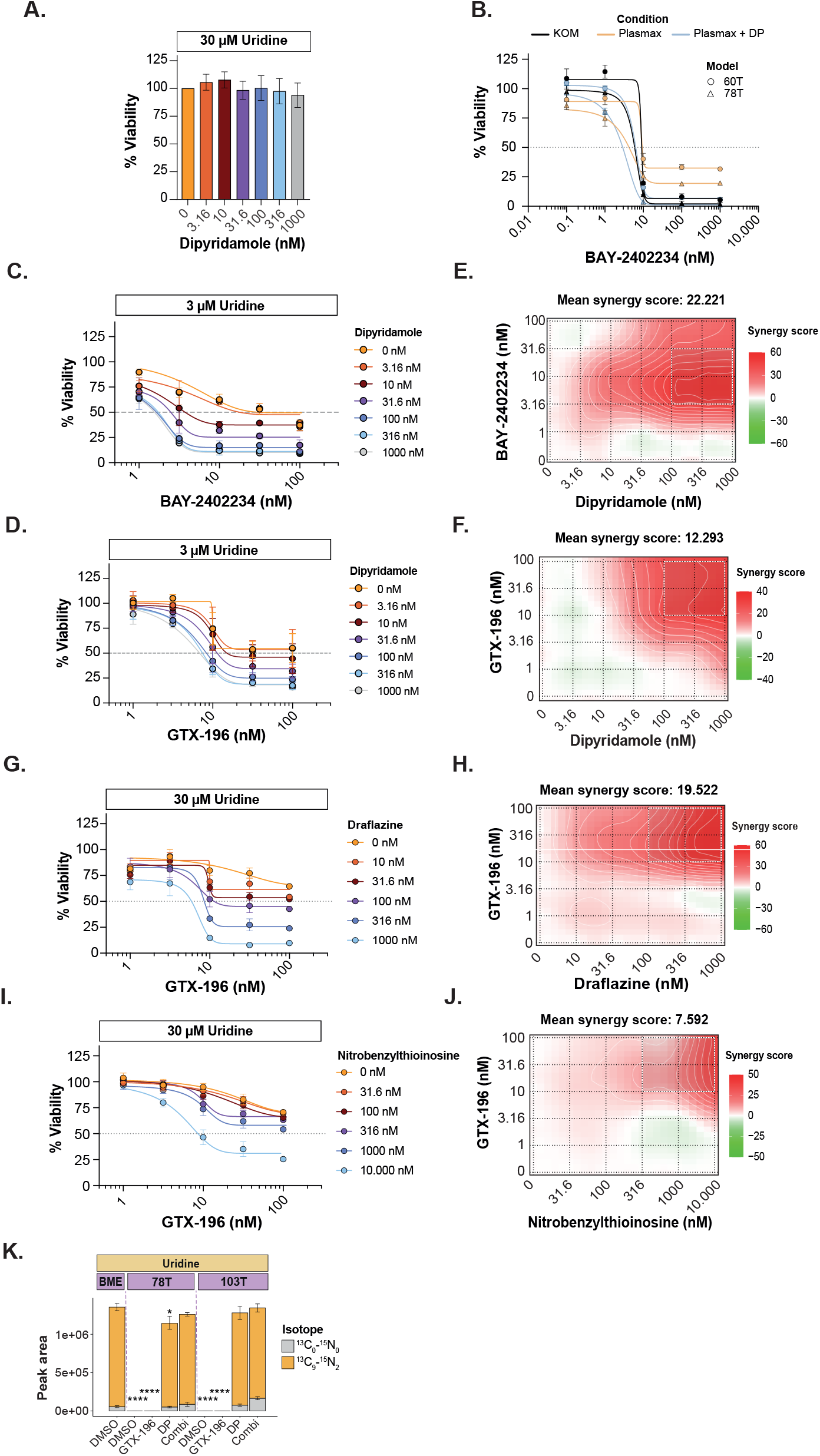
Combined targeting of DHODH and nucleotide salvage exerts synergistic anti-tumor effects in ecMRT under (supra-)physiologic nutrient levels. **A)** Bar graph depicting the average viability (%) of ecMRT tumoroids cultured in KOM supplemented with 30 μM uridine following 120-hour treatment with various concentrations of dipyridamole. Data represent the mean ± SD of *n = 3* independent experiments, each consisting of technical triplicates. Viability values were normalized to the DMSO vehicle control (100%). **B)** Dose-response curves for ecMRT tumoroid models 60T and 78T treated with GTX-196 for 120 hours in Plasmax™ medium, with or without the hENT1/2 inhibitor dipyridamole (500 nM). ecMRT tumoroids treated with GTX-196 in KOM medium served as a reference. Data represent mean ± SD from three independent experiments, each performed with technical duplicates. Data are normalized to DMSO vehicle in KOM (100%). The grey dotted horizontal line represents a viability of 50% (IC_50_). **C-D)** Dose-response curves for ecMRT tumoroid models 60T treated with BAY-2402234 (panel **C**) and 103T treated with GTX-196 (panel D) for 120 hours in KOM supplemented with 3 μM uridine, with or without different concentrations of hENT1/2 inhibitor dipyridamole. Data represent mean ± SD from three independent drug matrix experiments. Data are normalized to DMSO vehicle (100%). The grey dotted horizontal line represents a viability of 50% (IC_50_). **E-F)** Two-dimensional (2D) synergy landscapes visualized as contour plots, displaying ZIP (Zero Interaction Potency) synergy scores^30^ across concentration matrices for BAY-2402234 combined with dipyridamole (panel **E**) and GTX-196 combined with dipyridamole (panel **F**) (corresponding to figure panels **C** and **D**). ZIP scores >10 indicate synergistic interactions, scores between -10 and 10 represent additive effects, and scores <–10 indicate antagonism. Regions outlined in white represent the most synergistic concentration combinations. Synergy landscapes were generated using SynergyFinder software^31^. **G**,**I)** Dose-response curves for ecMRT tumoroid model 103T treated with GTX-196 for 120 hours in KOM supplemented with 30 μM uridine, with or without different concentrations of hENT inhibitors draflazine (panel **G**) or nitrobenzylthioinosine (NBMPR; panel I). Data represent mean ± SD from three independent drug matrix experiments. Data are normalized to DMSO vehicle (100%). The grey dotted horizontal line represents a viability of 50% (IC_50_). **H**,**J**) Two-dimensional (2D) synergy landscapes visualized as contour plots, displaying ZIP (Zero Interaction Potency) synergy scores^30^ across concentration matrices for GTX-196 combined with draflazine (panel **H**) and GTX-196 combined with NBMPR (panel **J**) (corresponding to figure panels **G** and **I**). ZIP scores >10 indicate synergistic interactions, scores between -10 and 10 represent additive effects, and scores <–10 indicate antagonism. Regions outlined in white represent the most synergistic concentration combinations. Synergy landscapes were generated using SynergyFinder software^31^. **K)** Isotopic labeling pattern of extracellular uridine levels measured in the medium of BME (no cells) and two different ecMRT tumoroids following 24-hour incubation with 3⍰μM or 30⍰μM [^13^C_9_-^15^N_2_]-uridine upon 48-hour treatment with DMSO vehicle, 5 nM GTX-196, 10 μM DP, or the combination thereof. All conditions were statistically compared to the BME (no cells) reference group. (*, *p* < 0.05; **, *p* < 0.01; ***, *p* < 0.001; ****, *p* < 0.0001).

**Table S1.**
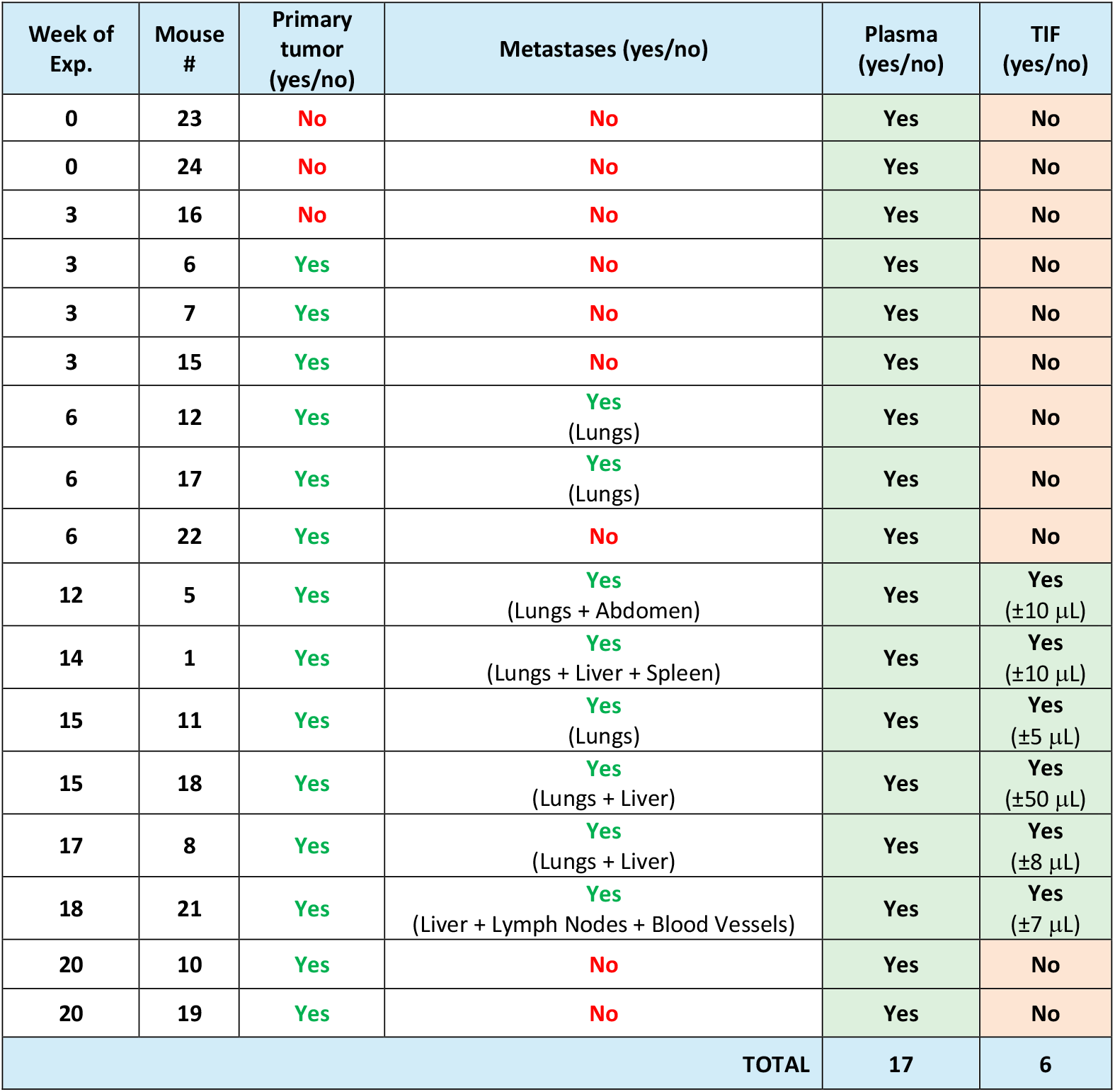
Summary of mouse cohort, tumor staging, and sample collection details.

| Week of Exp. | Mouse # | Primary tumor (yes/no) | Metastases (yes/no) | Plasma (yes/no) | TIF (yes/no) |
| --- | --- | --- | --- | --- | --- |
| 0 | 23 | No | No | Yes | No |
| 0 | 24 | No | No | Yes | No |
| 3 | 16 | No | No | Yes | No |
| 3 | 6 | Yes | No | Yes | No |
| 3 | 7 | Yes | No | Yes | No |
| 3 | 15 | Yes | No | Yes | No |
| 6 | 12 | Yes | Yes<br>(Lungs) | Yes | No |
| 6 | 17 | Yes | Yes<br>(Lungs) | Yes | No |
| 6 | 22 | Yes | No | Yes | No |
| 12 | 5 | Yes | Yes<br>(Lungs + Abdomen) | Yes | Yes<br>(±10 µL) |
| 14 | 1 | Yes | Yes<br>(Lungs + Liver + Spleen) | Yes | Yes<br>(±10 µL) |
| 15 | 11 | Yes | Yes<br>(Lungs) | Yes | Yes<br>(±5 µL) |
| 15 | 18 | Yes | Yes<br>(Lungs + Liver) | Yes | Yes<br>(±50 µL) |
| 17 | 8 | Yes | Yes<br>(Lungs + Liver) | Yes | Yes<br>(±8 µL) |
| 18 | 21 | Yes | Yes<br>(Liver + Lymph Nodes + Blood Vessels) | Yes | Yes<br>(±7 µL) |
| 20 | 10 | Yes | No | Yes | No |
| 20 | 19 | Yes | No | Yes | No |
| TOTAL |  |  |  | 17 | 6 |

**Table S2.** Proxy labeled metabolites used for normalization of unlabeled metabolites when direct labeled matches were unavailable.

| 12C_Metabolite | Isotopologue | Formula | RT (min) | m/z | Polarity | 13C.yeast_Proxy | Isotopologue | Formula | RT (min) | m/z | Polarity |
| --- | --- | --- | --- | --- | --- | --- | --- | --- | --- | --- | --- |
| Uridine | [M+0] | C9H12N2O6 | 7,4 | 243,0659 | M-H | Thymidine | [M+10] | [13]C10H14N2O5 | 5,48 | 251,1165 | M-H |
| CMP | [M+0] | C9H14N3O8P | 17,23 | 322,0446 | M-H | UMP | [M+9] | [13]C9H13N2O9P | 16,42 | 332,0588 | M-H |
| Hypoxanthine | [M+0] | C5H4N4O | 7,8 | 137,045 | M+H | Adenine | [M+5] | [13]C5H5N5 | 6,77 | 141,079 | M+H |
| Uracil | [M+0] | C4H4N2O2 | 7,42 | 111,0199 | M-H | Thymidine | [M+10] | [13]C10H14N2O5 | 5,48 | 251,1165 | M-H |
| UTP | [M+0] | C9H15N2O15P3 | 20,13 | 483,9831 | M-H | UMP | [M+9] | [13]C9H13N2O9P | 16,42 | 332,0588 | M-H |
| CDP | [M+0] | C9H15N3O11P2 | 19,12 | 403,005 | M-H | UMP | [M+9] | [13]C9H13N2O9P | 16,42 | 332,0588 | M-H |
| Inosine | [M+0] | C10H12N4O5 | 9,23 | 269,0888 | M+H | Adenosine | [M+10] | [13]C10H13N5O4 | 6,37 | 278,1376 | M+H |
| cAMP | [M+0] | C10H12N5O6P | 8,25 | 328,0452 | M-H | AMP | [M+10] | [13]C10H14N5O7P | 14,86 | 356,09 | M-H |
| GMP | [M+0] | C10H14N5O8P | 18,12 | 362,0502 | M-H | AMP | [M+10] | [13]C10H14N5O7P | 14,86 | 356,09 | M-H |
| ATP | [M+0] | C10H16N5O13P3 | 17,66 | 506,9870 | M-H | ADP | [M+10] | [13]C10H15N5O10P2 | 16,52 | 437,0631 | M-H |
| Dimethyllysine | [M+0] | C8H18N2O2 | 21,05 | 174,1444 | M+H | LYS | [M+6] | [13]C6H14N2O2 | 23,99 | 153,133 | M+H |
| ADMA | [M+0] | C8H18N4O2 | 21,1 | 203,1509 | M+H | ARG | [M+6] | [13]C6H14N4O2 | 24,93 | 181,1391 | M+H |
| NMMA | [M+0] | C7H16N4O2 | 23,44 | 189,1353 | M+H | ARG | [M+6] | [13]C6H14N4O2 | 24,93 | 181,1391 | M+H |
| TYR | [M+0] | C9H11NO3 | 12,78 | 180,0655 | M+H | PHE | [M+9] | [13]C9H11NO2 | 7,98 | 175,1166 | M+H |
| TRP | [M+0] | C11H12N2O2 | 10,77 | 205,0972 | M+H | PHE | [M+9] | [13]C9H11NO2 | 7,98 | 175,1166 | M+H |
| GLY | [M+0] | C2H5NO2 | 16,13 | 76,0393 | M+H | SER | [M+3] | [13]C3H7NO3 | 15,65 | 109,0599 | M+H |
| Dimethylglycine | [M+0] | C4H9NO2 | 11,42 | 104,0706 | M+H | SER | [M+3] | [13]C3H7NO3 | 15,65 | 109,0599 | M+H |
| Homocysteine | [M+0] | C4H9NO2S | 12,92 | 136,0426 | M+H | SER | [M+3] | [13]C3H7NO3 | 15,65 | 109,0599 | M+H |
| Betaine | [M+0] | C5H11NO2 | 9,94 | 118,0863 | M+H | L-Carnitine | [M+7] | [13]C7H15NO3 | 12,95 | 169,136 | M+H |
| Citrate | [M+0] | C6H8O7 | 20,66 | 191,0197 | M-H | Cis-Aconitate | [M+6] | [13]C6H6O6 | 19,21 | 179,029 | M-H |
| Methylnicotinamide | [M+0] | C7H8N2O | 23,39 | 137,0712 | M+H | Niacinamide | [M+6] | [13]C6H6N2O | 5,47 | 129,0756 | M+H |

